# Morphological and heat-tolerance traits are associated with progression and impact of, but not vulnerability to, tree decline

**DOI:** 10.1101/2025.09.25.677476

**Authors:** Sabina M. Aitken, Pieter A. Arnold, Matthew T. Brookhouse, Alicia M. Cook, Lisa M. Danzey, Rosalie J. Harris, Andy Leigh, Adrienne B. Nicotra

**Author notes:** **Corresponding author:** Adrienne Nicotra. **Author contributions (CRediT)** SA led and was involved with all aspects of the work as were AN and MB. PA, AC, LD, RH and AL contributed to data analysis, visualisation. All authors contributed to interpretation, reviewing and editing the manuscript.

## Abstract

Warming and drying climate trends are driving tree dieback worldwide with broad-reaching impacts on ecosystem services. Studying decline is unavoidably a retrospective exercise in which researchers are challenged to determine whether trait values that are associated with dieback are drivers versus consquences of decline. In this study, we use the subalpine snow gum (*Eucalyptus pauciflora, ssp niphophila*) as a case study to illustrate how to identify whether plant traits may explain vulnerability of individual trees, assess how progression of dieback symptoms affect traits and physiological tolerance, and ask whether those changes could exacerbate decline. While the impact of drought on tree mortality has been well documented, we consider the potential role of heat which has received considerably less attention. We assessed changes in leaf and stem morphology, and stomatal anatomy across a dieback severity and an abiotic elevation gradient. We also assessed the relationship between these traits, photosystem heat tolerance and dieback progression, using results to model leaf viability under current and future climate scenarios. While severely symptomatic trees exibited trait values indicating water stress, trees with low or moderate dieback were not different from unaffected ones. Thus the differences in severely affected trees are likely responses to water stress caused by woodborer girdling and provide no evidence of underlying trait difference driving vulnerability. Severely symptomatic trees, however, had lower photosystem heat tolerance and models indicated leaves were likely to accumulate lethal damage to photosystems within a growing season, thus contributing to a feedback cycle of decline even under current thermal regimes.

**Highlights:**

- Accounting for abiotic, spatial variation explains complex relationships in dieback research
- Insect mediated dieback induces trait variation consistent with water limitation
- No evidence of variation in vulnerability to dieback among snow gum individuals
- Dieback-induced reduction in heat tolerance compounds heat load effects
- Warming induced heat loading will compromise carbon gain and exacerbate decline

## 1. Introduction

Recent warming and drying trends have been linked to increases in tree and stand mortality across forested biomes globally (Adams et al., 2009; Allen et al., 2010; Andrus et al., 2024; Hammond et al., 2022; Hartmann et al., 2022). These dieback events have the potential to significantly alter biodiversity and ecosystem services. Already, tree dieback has been associated with declines in co-occurring species, changes to water catchments and streamflow, and reduced carbon sequestration (Boyd et al., 2013; Camarero et al., 2015; Martin et al., 2015; Mitchell et al., 2014). Dieback events are complex, typically reflecting a combination of abiotic stressors (e.g. climatic or edaphic extremes) and altered biotic interactions (e.g. insects or pathogens) that together bring about tree decline (Jurskis, 2005; Manion, 1991; Mueller-Dombois, 1988).

Variation in plant traits along environmental gradients may correlate with dieback susceptibility and assist to explain spatial patterns in dieback distribution. Variation in traits may increase or decrease tolerance of climatic stressors or contribute to vulnerability to insects or pathogens. For example, when faced with reduced water availability, traits associated with plant-water use and drought stress may contribute to vulnerability of individual trees (Choat et al., 2018; McDowell et al., 2008). There is also the possibility however, for variation in plant traits to occur as a result of the onset of dieback, not associated with trait-related vulnerability. Given that trees are long lived organisms and dieback research is most often conducted in the field and after onset of the event, disentangling complex environmental variation is challenging. It is important to isolate trait variation relationships with dieback severity independently of trends associated with environmental gradients such as elevation. For example, key morphological traits such as leaf area, leaf mass per unit area (LMA), and Huber value (ratio of stem cross-sectional area to distal leaf area) generally exhibit a conservative response to water stress induced by abiotic (e.g. drought) or biotic (e.g. stem girdling borers) drivers. These include smaller leaf area, lower Huber values, higher LMA and reduced stomatal size and density, which limit water loss (Carter & White, 2009; Hosseini et al., 2019; Lin et al., 2021; Niinemets, 2001; Poorter et al., 2009; Wright et al., 2017). In many cases these same conservative leaf traits are also associated with abiotic factors, for example, higher elevation where cooler temperatures limit leaf expansion and favour thicker, more frost resistant leaves.

As dieback progresses, physiological tolerance, such as photosystem heat tolerance, may also be affected by and potentially contribute to tree-level decline. Water limitation may result in increased heat tolerance due to acclimation to limited evaporative cooling and associated higher leaf temperatures (Cook et al., 2021; Havaux, 1992; Valladares & Pearcy, 1997). Alternatively, functionality may decline under drought conditions as the activation of multiple stress-response pathways limit resources available for photosystem function (Ben Rejeb et al., 2014; Zhu et al., 2021). The duration of exposure to heat also affects photosystem function, whereby the temperature that the leaf tissue can functionally tolerate decreases as duration of heat exposure increases, i.e., with increased heat load (Cook et al., 2024; Faber et al., 2024; Neuner & Buchner, 2023). It is unclear whether biotic stressors associated with dieback would enhance or diminish photosystem heat stress tolerance and further affect the trees’ vulnerability to warming environmental conditions. While the importance of water limitation to global dieback events is well understood, the question of whether temperature could be a direct driver of decline has received less attention (Allen et al., 2010; Anderegg et al., 2015; Camarero et al., 2015; Choat et al., 2018; Meir et al., 2015).

Sub-alpine and montane ecosystems may be particularly vulnerable to climate change-induced dieback as rates of warming at high elevations exceed the global average (Gobiet et al., 2014; IPCC, 2022). As temperatures increase, reduced thermal constraints on insects and pathogenic organisms at high elevations may increase the susceptibility of these forests to biotic infestations (Anderegg et al., 2015; Raffa et al., 2008). For example, warming winter temperatures have increased over-winter survival and reproductive rates of bark beetles, contributing to widespread decline of *Pinus*-dominated forests in North America and Europe (Adams et al., 2009; Jaime et al., 2024; Mitton & Ferrenberg, 2012; Raffa et al., 2008). Causes and consequences of decline in evergreen broadleaf forests or in southern hemisphere forests more generally has not received comparable attention, despite increasingly alarming reports of forest decline (Anderegg et al., 2012; Ross & Brack, 2015). One such example is the decline occurring in snow gum in Australia (*Eucalyptus pauciflora* ssp. *niphophila* (Maiden & Blakely) L.A.S. Johnson & Blaxell) that has emerged in the past decade. This decline appears to be aligned with the effect of warmer temperatures and drought on the trees which is, like the northern hemisphere examples, mediated by a bark boring beetle, in this case from the globally significant *Phoracantha* group: *P. mastersi*.

Using the current dieback event in snow gum as a case study, we address two global knowledge gaps around forest decline. First, we demonstrate an effective approach to disentangle the impacts of plant-traits from potetial abiotic drivers (elevation) of dieback response given the retrospective nature of dieback research. We postulate that if certain leaf trait values are associated with vulnerability to dieback, then any tree showing onset of dieback will exhibit similar trait values, whether moderately or severely affected. Whereas, if trees with low dieback severity resemble those with no signs of decline, then trait differences are unlikely to represent variation in vulnerability. Second, we provide an important example of feedback between dieback severity and declining physiological tolerance, specifically photosystem thermal tolerance. Past studies suggest that heat tolerance may increase or decrease with water stress (e.g. associated with drought and bark damage) (Cook et al., 2021; Havaux, 1992; Valladares & Pearcy, 1997; Zhu et al., 2021), however, the relative impact of thermal stress on instigating tree mortality has not been examined. Here we apply dynamic models of heat load to explore the influence of exposure duration on leaf heat tolerance and its implications for leaf viability under current and future climate scenarios (i.e. thermal-load sensitivity (Arnold et al., 2025)). We examined whether water limitation would improve or reduce heat tolerance and predicted that in severely affected individuals heat tolerance would be reduced such that heat load models would highlight a positive feedback cycle that exacerbates tree decline in snow gum and other dieback events elsewhere.

## 2. Materials and Methods

### 2.1 Study species and field site

This study was conducted in subalpine woodlands in the Snowy River valley between 1680 and 1890 m asl in Kosciuszko National Park (KNP; Figure 1, see also supplemental information, S1), New South Wales, Australia. Consistent with forest composition throughout KNP above 1600 m asl, stands within the study area are comprised entirely of sparsely spaced, multi-stemmed *E. pauciflora* ssp. *niphophila* with a shrub or grassy understory.

**Figure 1.**
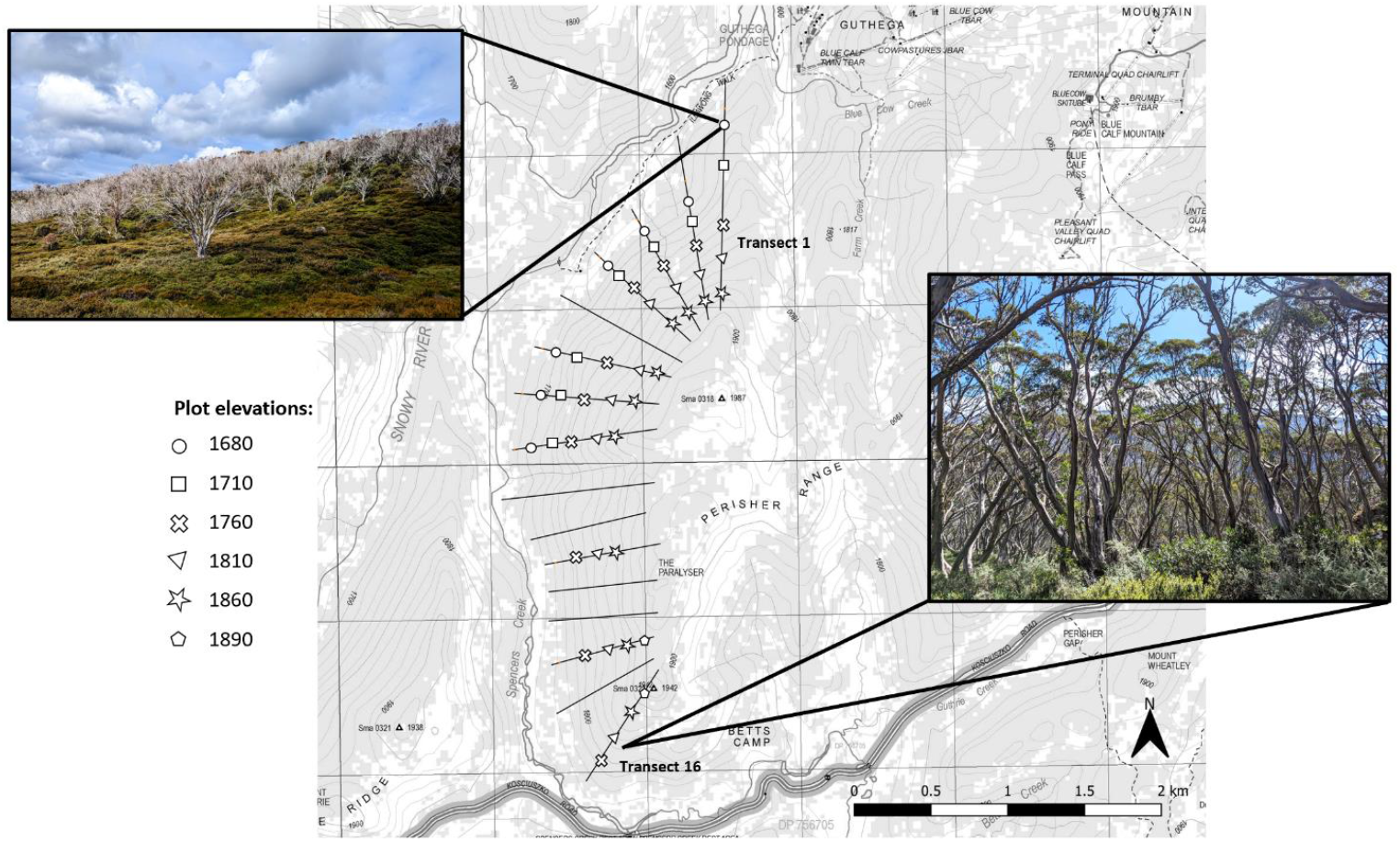
Location of 46 study plots, along 10 transects on slopes of The Paralyser, Kosciuszko National Park, NSW, Australia. Inset images show comparison of dieback severity at transect #1 (severe dieback) to transect #16 (largely unaffected by dieback). Study plots range in elevation from 1680 m asl to 1890 m asl.

Snow gum dieback was first reported in the late twentieth century as localised patch-scale decline (Banks, 1982, 1989; Shields, 1993), but is now widespread throughout the Australian Alps. Dieback affected trees show canopy decline as well as tissue damage distinctive of the native wood-boring beetle, *Phoracantha mastersi* (Pascoe; Cerambycidae). Other species of the genus *Phoracantha* have been documented to cause significant damage in *Eucalyptus* forestry plantations in Australia and overseas. Studies of these species note that the survival of the wood-boring larvae is aided by dry conditions, allowing larvae to reach cambial tissue more successfully (Da Conceição Caldeira et al., 2001; Hanks et al., 1991, 1999; Seaton et al., 2015). Larvae of *P. mastersi* feed circumferentially on cambial tissue of snow gum, consuming both the outer xylem and inner phloem tissue, effectively ring-barking the stem (Brookhouse et al., 2024). The presence and frequency of *P. mastersi* infestations is inversely correlated with elevation above 1600 m asl, the lower elevational limit of *E. pauciflora* ssp. *niphophila* (Brookhouse et al., 2024; Bryant et al., 2024). This elevational pattern raises two possibilities that require further exploration. First, thermal limitation of *P. mastersi* behaviour may suppress infestation rates at high elevation, consistent with outbreaks of insects in alpine and montane forests elsewhere (Bentz et al., 2010; Jaime et al., 2024). Second, temperature-mediated variation in plant traits may promote vulnerability to infestations at lower elevation.

To disentangle the role of elevation in dieback we established a network of temporary plots established on elevational transects positioned along a naturally occurring dieback gradient between the Guthega ski resort and Spencers Creek snow-depth monitoring station (36°24’31”S 148°21’28”E, Figure 1). Prior field surveys indicated that *P. mastersi* infestations declined in severity with increasing distance from Guthega ski resort as well as with increasing elevation (Brookhouse et al., 2024). Sixteen elevational transects were established to span the full range of forest cover within the study area, ranging from 1680 m to 1890 m asl. Variable-radius plots were established at six fixed elevations (1680, 1710, 1760, 1810, 1860 and 1890 m asl), using a basal area factor of 2. Consistent with Brookhouse et al. (2024), canopy health and level of wood-borer damage, were used to classify the severity of dieback of each tree as either none, low, moderate or severe (see S1a-c, and Figures S1 and S2 for more detail).

### 2.2 Effects of dieback severity and elevation on leaf morphology and stomatal anatomy

In each of 46 plots from a subset of ten of the original transects, three representative trees with live canopy were selected for sampling. Collection took place during the austral summer-autumn (February-April) of 2023-24. For leaf-trait measurements, small branches growing in full sun were retrieved from the upper canopy of the tree using an arborist’s throwline. Branches were wrapped in damp paper towel and stored in zip-lock bags in the field to prevent loss of leaf water then measured at the field laboratory within ∼8 hours of collection.

Becuase spread of dieback in the study area was quite recent, symptoms of dieback may have begun in the previous 12 months. To test for progression of dieback symptoms, leaves from both the current and previous growing season were sampled. For each tree, three healthy leaves that expanded during the current growing season (2022/2023) (hereafter referred to as ‘young leaves’) and three healthy leaves that expanded during the preceding growing season (2021/2022; hereafter ‘old leaves’) were randomly sampled from collected branches for leaf trait measurements. The distinction between young and old leaves was made visually based on location on the branch and leaf and petiole colour (see S1d, Figure S3). Leaf area, leaf mass per unit area (LMA) were measured for each leaf, stomatal size, stomatal density and stomatal pore index (Lin et al., 2021; Sack et al., 2003) were measured on one leaf per leaf age class per tree and Huber values for one branch per tree (see supplemental information S1e-f).

Given hydraulic damage caused by woodborers, we predicted that dieback affected trees would show conservative leaf traits, generally associated with water stress (including smaller leaf areas, stomatal size, stomatal density and Huber values). We further anticipated that the most recent cohort of leaves would show stronger signals of dieback than the previous cohort.

### 2.3 Photosystem heat tolerance

Leaves for photosystem heat tolerance assays were sampled from *E. pauciflora* ssp. *niphophila* trees at a single site (36°24’35.9”S 148°24’52.2”E, 1780 m asl), independent of the transect network used for leaf-trait sampling. This site, which falls in the mid-elevational range of dieback-affected trees, was selected because it contained a range of dieback severity. Leaf samples were collected from eight trees within each of three dieback-severity classes representing none, moderate, and severe symptoms classified as described above.

We measured both a point and a cumulative measure of photosystem heat tolerance. As a point metric, we measured the critical thermal threshold (T_crit_) at which basal chlorophyll fluorescence (F_0_) begins to rise rapidly, indicative of the temperature at which irreversible photosystem damage occurs (Arnold et al., 2021; Schreiber & Berry, 1977). Because T_crit_ does not account for the influence of duration on thermal stress, we also considered thermal-load sensitivity (TLS), using maximum quantum yield of PSII (Fv/Fm) response to steady-state heat assay/challenge. This TLS analysis offered insight into the effect of exposure duration in determining the impact of temperature stress on plant physiological processes (Arnold et al., 2025; Cook et al., 2024; Faber et al., 2024; Neuner & Buchner, 2023). For detailed methods see supplemental information (S2).

### 2.4 Simulating thermal load sensitivity effects under field thermal regimes

Finally, as a simulated demonstration of the impact of accumulating heat load to which the leaves were exposed in the 2022-23 summer, we modelled realistic local field leaf temperatures using the microclimate and ectotherm models of ‘NicheMapR’ R package (Kearney & Leigh, 2024; Kearney & Porter, 2017, 2020). We simulated microclimate at 2 m above ground level using the *micro_era5* function (Kearney et al., 2020; Klinges et al., 2022), which extracts climatic and topographic variables from the ERA5 dataset (Hersbach et al., 2020) at the study location from which TLS leaves were sampled (36°24’35.9”S 148°24’52.2”E, 1780 m asl). Leaves were specified to be an ellipsoid shape (length = 66.7 mm, width = 18.4 mm and thickness = 0.63 mm) with tissue density = 1 g cm^-3^.

Previous field measurements of *E. pauciflora* ssp. *niphophila* were used to parameterise leaf shape and other functional traits required for the modelling: leaf wet weight = 0.53 g (mean summer wet weight) and solar absorptivity of the leaf = 0.93. Plausible values of mean leaf diffusive conductance = 0.30 mol m^-2^ s^-1^, and maximum leaf diffusive conductance = 0.35 mol m^-2^ s^-1^ were derived from relevant elevation populations of *E. pauciflora* from Körner & Cochrane (1985). Leaf temperature models (derived using leaf mode in the *ectotherm* function) were then run for two stomatal states: open during appropriate diurnal conditions or always closed, representing functional and water-stressed leaves, respectively. Using the TLS parameter estimates (CT_max_′ and *z*) from none, moderate, and severe dieback severity classes measured in this study, we fit ‘thermal tolerance landscape’ models (Rezende et al., 2014, 2020). These models estimate the leaf survival probability (%) at each day, based on the temperatures that the leaves reach across time (heat load), and their capacity to tolerate heat load before accumulating damage to PSII. We then fit an +4°C increase in temperature over the same time course to assess the extent to which heat load would lead to compounded leaf damage under a high climate-warming scenario (SSP3, (IPCC, 2022), for open and closed stomata states, and different dieback-severity classes.

### 2.5 Data analysis

All statistical analyses were performed using R Statistical Software, v4.2.3 (R Core Team, 2021). For transect-based data (leaf morphology and stomatal traits), the ‘nlme’ R package (Pinheiro et al., 2023) was used to fit linear mixed-effects regression (LMER) models. These models assessed the change in leaf traits associated with dieback severity (none, low, moderate, severe), elevation and leaf age (young, old) as fixed effects. The identity of each sample tree was included as a random factor where multiple measurements were taken from a single tree. All models also included a spatial correlation structure (derived from the position of each sample tree) to account for potential spatial autocorrelation in dieback severity. Models were compared using corrected Akaike Information Criterion (AICc) and for all traits except LMA final models did not include interactions (see supplemental information S3). To analyse heat tolerance data, we fit similar LMER models, but without elevation and spatial autocorrelation terms as these data were collected at a single site. Dieback severity and leaf age were fixed effects while tree and sampling day were random variables.

## 3. Results

### 3.1 Effects of elevation and dieback severity on leaf and stem traits

Contrary to our expectations, leaf morphological traits did not vary significantly with elevation. Further, while trait variation between dieback-severity classes broadly aligned with our predictions (Table 1), that variation largely only distinguished leaves from trees with the highest severity rating from those with no, low or moderate dieback severity. Thus, our results do not indicate any inherent differences in leaf traits that might predispose these trees to dieback.

**Table 1.** Effects of dieback, elevation and leaf age on leaf morphology and stomata anatomy of Eucalyptus pauciflora ssp. niphophila: results from linear mixed effects regression models. Note uber value was measured at the branch level so there is no effect of leaf age for this trait. Intercept corresponds to old leaves from unaffected trees.

|  | Leaf area<br>(cm<sup>2</sup>)<br>n=748* |  |  | LMA<br>(g/m<sup>2</sup>)<br>n=748 |  |  | Huber value<br>(cm<sup>2</sup>/m<sup>2</sup>)<br>n=114 |  |  | Stomata length<br>(µm)<br>n=233 |  |  | Stomata density<br>(no. stomata/mm<sup>2</sup>)<br>n=233 |  |  | Stomatal pore index<br>n=233 |  |  |
| --- | --- | --- | --- | --- | --- | --- | --- | --- | --- | --- | --- | --- | --- | --- | --- | --- | --- | --- |
|  | Est. | (se) | P-value | Est. | (se) | P-value | Est. | (se) | P-value | Est. | (se) | P-value | Est. | (se) | P-value | Est. | (se) | P-value |
| Intercept | 2.181 | 0.547 | <0.001 | 397.46 | 70.049 | <0.001 | 18.551 | 9.436 | 0.052 | 44.451 | 10.216 | <0.001 | 4.142 | 0.509 | <0.001 | 2.055 | 0.653 | 0.002 |
| Elevation | 9.5e-5 | 3.0e-4 | 0.752 | -0.024 | 0.038 | 0.538 | -0.007 | 0.005 | 0.183 | -0.008 | 0.006 | 0.148 | 4.79e-4 | 2.82e-4 | 0.091 | 2.7e-4 | 3.60e-4 | 0.453 |
| Dieback (low) | -0.037 | 0.040 | 0.358 | -9.134 | 5.733 | 0.114 | -0.592 | 0.655 | 0.368 | 0.411 | 0.625 | 0.511 | -0.047 | 0.026 | 0.066 | -0.008 | 0.0411 | 0.846 |
| Dieback (mod) | -0.047 | 0.060 | 0.431 | -6.981 | 8.534 | 0.415 | -0.334 | 0.945 | 0.724 | **-2.762** | **0.927** | **0.003** | 0.035 | 0.038 | 0.352 | -0.102 | 0.0597 | 0.090 |
| Dieback (severe) | **-0.198** | **0.056** | **0.001** | -9.680 | 7.841 | 0.219 | **2.772** | **0.929** | **0.004** | **-2.024** | **0.937** | **0.032** | 0.037 | 0.038 | 0.326 | -0.025 | 0.0610 | 0.686 |
| Leaf age (young) | **-0.235** | **0.018** | **<0.001** | **-99.33** | **3.152** | **<0.001** | - | - | - | -0.427 | 0.307 | 0.165 | **0.035** | **0.016** | **0.034** | 0.007 | 0.0231 | 0.772 |
| Db(low):L(young) | - | - | - | **18.174** | **4.964** | **<0.001** | - | - | - | - | - | - | - | - | - | - | - | - |
| Db(mod):L(young) | - | - | - | 8.165 | 7.454 | 0.273 | - | - | - | - | - | - | - | - | - | - | - | - |
| Db(sev):L(young) | - | - | - | **14.885** | **6.297** | **0.028** | - | - | - | - | - | - | - | - | - | - | - | - |
| Spatial correlation structure | *AICc with CorStruc:* -33.61<br>*AICc without:* 23.87 |  |  | *AICc with CorStruc:* 7059.7<br>*AICc without:* 7088.1 |  |  | *AICc with CorStruc:* 570.10<br>*AICc without:* 565.83 |  |  | *AICc with CorStruc:* 1230.9<br>*AICc without:* 1264.7 |  |  | *AICc with CorStruc:* -211.31<br>*AICc without:* -190.64 |  |  | *AICc with CorStruc:* -18.17<br>*AICc without:* -1.92 |  |  |
| Random effect of Tree ID | *Marginal R<sup>2</sup>:* 0.214<br>*Conditional R<sup>2</sup>:* 0.414 |  |  | *Marginal R<sup>2</sup>:* 0.674<br>*Conditional R<sup>2</sup>:* 0.739 |  |  | - |  |  | - |  |  | - |  |  | - |  |  |

\*Sample size n=748 corresponds to 128 trees × 2 leaf ages × 3 leaf replicates, see S1f for full account of sample size.

Across all dieback severities, old leaves (formed during 2021/22) were larger than young leaves (formed during 2022/23) (Figure 2A, Table 1). Trees with severe dieback had smaller leaves than those from less severely affected, being significantly smaller than unaffected trees across both leaf age cohorts (Figure 2A, Table 1). Average leaf area for trees in low and moderate dieback categories did not differ from that of unaffected trees (Table 1). In addition to smaller leaves, severely affected trees also had significantly less leaf area per given stem than unaffected stems so that Huber values were on average 60% greater in severely affected trees (Figure 2C, Table 1).

**Figure 2.**
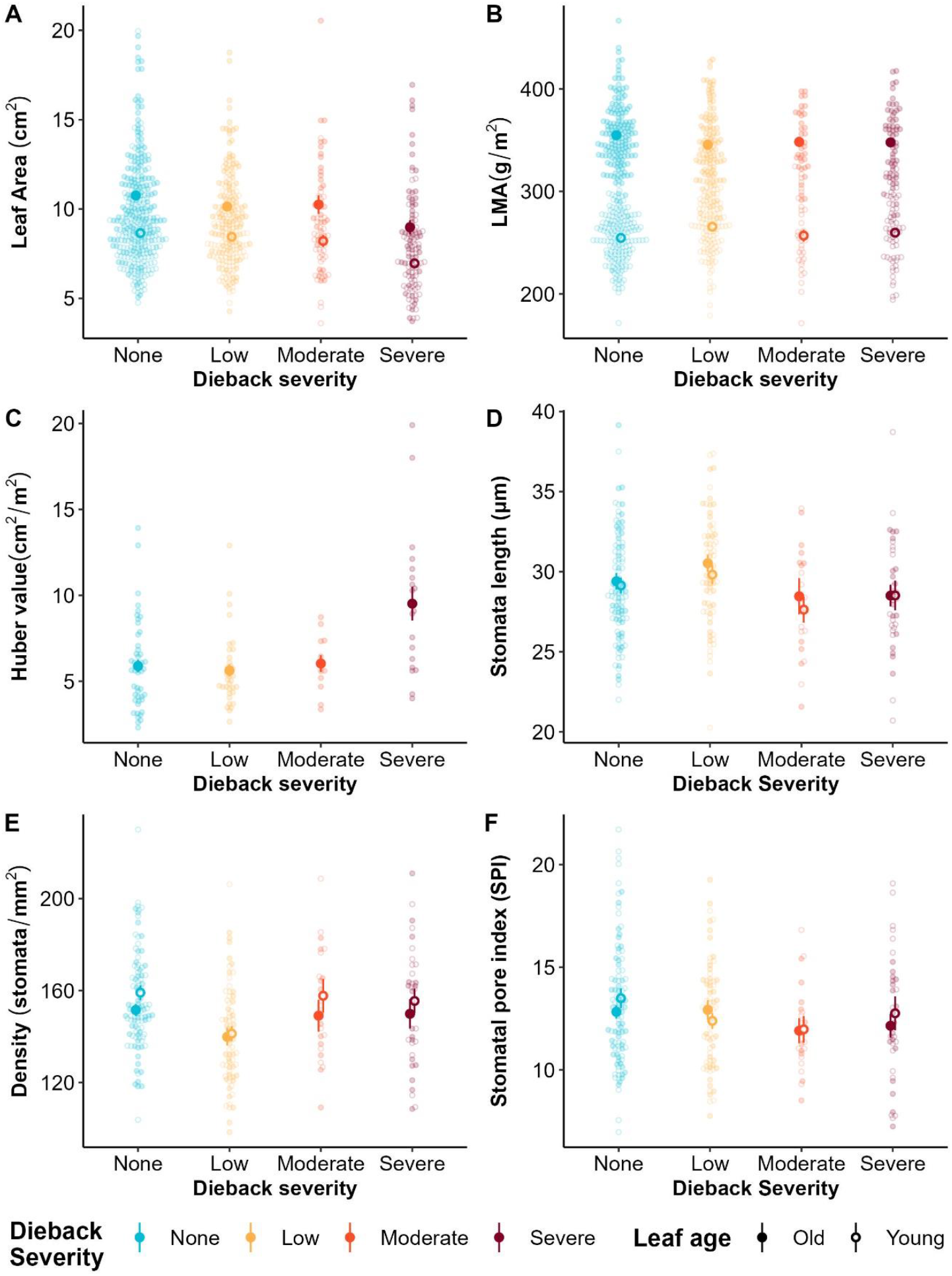
The response of morphological and stomatal traits to increasing dieback severity in *Eucalyptus pauciflora* ssp. *niphophila*. **(A)** individual leaf area (cm^2^), **(B)** leaf mass per area (LMA, g m^-2^) and **(C)** Huber value (ratio of stem cross-sectional area (cm^2^) to distal leaf area (m^2^)), **(D)** stomatal length (µm), **(E)** stomatal density (no. stomata mm^-2^) and **(F)** stomatal pore index (SPI, dimensionless), a proxy for maximum conductance potential considering both stomatal density and stomatal length. Huber value was measured on a branch level so there is no effect of leaf age. Closed circles indicate ‘old’ leaves i.e. those that expanded in the 2021/2022 growing season, while open circles indicate ‘young’ leaves i.e. those that expanded in the 2022/2023 growing season. Error bars reflect standard error about the mean.

There was no change in LMA with increasing dieback severity (Figure 2B); however, young leaves had lower LMA than old leaves (Figure 2B). The model indicated a significant interaction between dieback severity and leaf age because the extent of difference in LMA between young and old leaves varied among dieback severity classes.

We did not detect any decrease in stomatal size or density with elevation, however, as predicted, moderately and severely dieback-affected trees had significantly smaller stomata than those without evidence of dieback (Figure 2D, Table 1). Young leaves had significantly higher stomatal density than old leaves (Table 1) but there was no change in stomatal density with increasing dieback severity (Figure 2E). While the average stomatal pore index (SPI) of moderate and severely dieback-affected trees was slightly lower than that of unaffected trees, the difference was not statistically significant (Figure 2F, Table 1).

### 3.2 Photosystem heat tolerance

The maximum quantum yield of PSII (F_V_/F_M_) measured prior to temperature ramping was significantly lower in trees with severe dieback than unaffected trees, indicating reduced photosystem health associated with advanced *P. mastersi* infestation (Figure 3A, Table 2). The upper critical temperature at which stress is incurred within PSII, T_crit_, was also significantly lower in severely affected than unaffected trees (Figure 3B, Table 2). There was a decline of ∼1°C in T_crit_ with each increasing level of dieback severity (Figure 3B) and trees with higher initial F_V_/F_M_ generally had a higher T_crit_ (see Figure S5). There was no effect of leaf age on initial F_V_/F_M_ or T_crit_ values (Table 2).

**Table 2.**
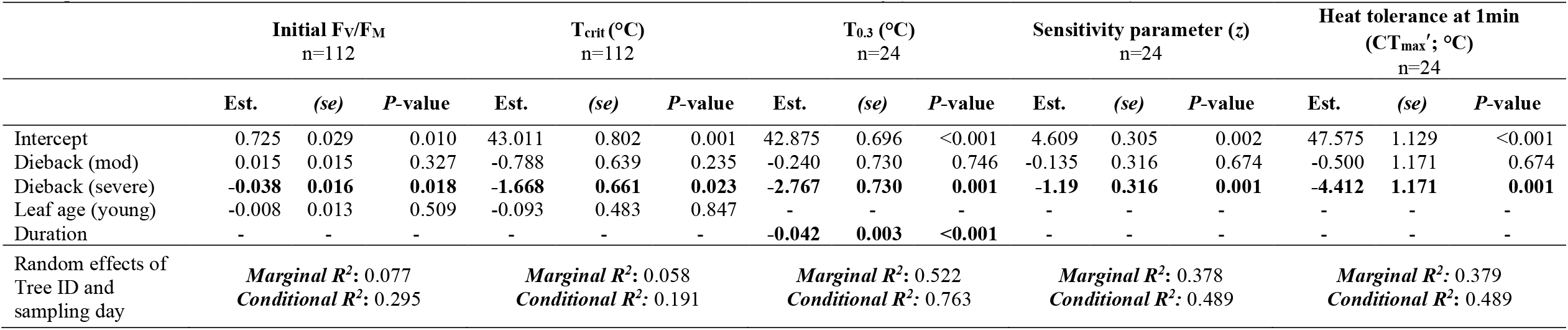
Effects of dieback and leaf age on photosystem heat tolerance measures in *Eucalyptus pauciflora* ssp. *niphophila*: results from linear mixed effects regression models. Intercept orresponds to old leaves from unaffected trees. Heat tolerance data included 3 classes of dieback severity (none, moderate, and severe).

**Figure 3.**
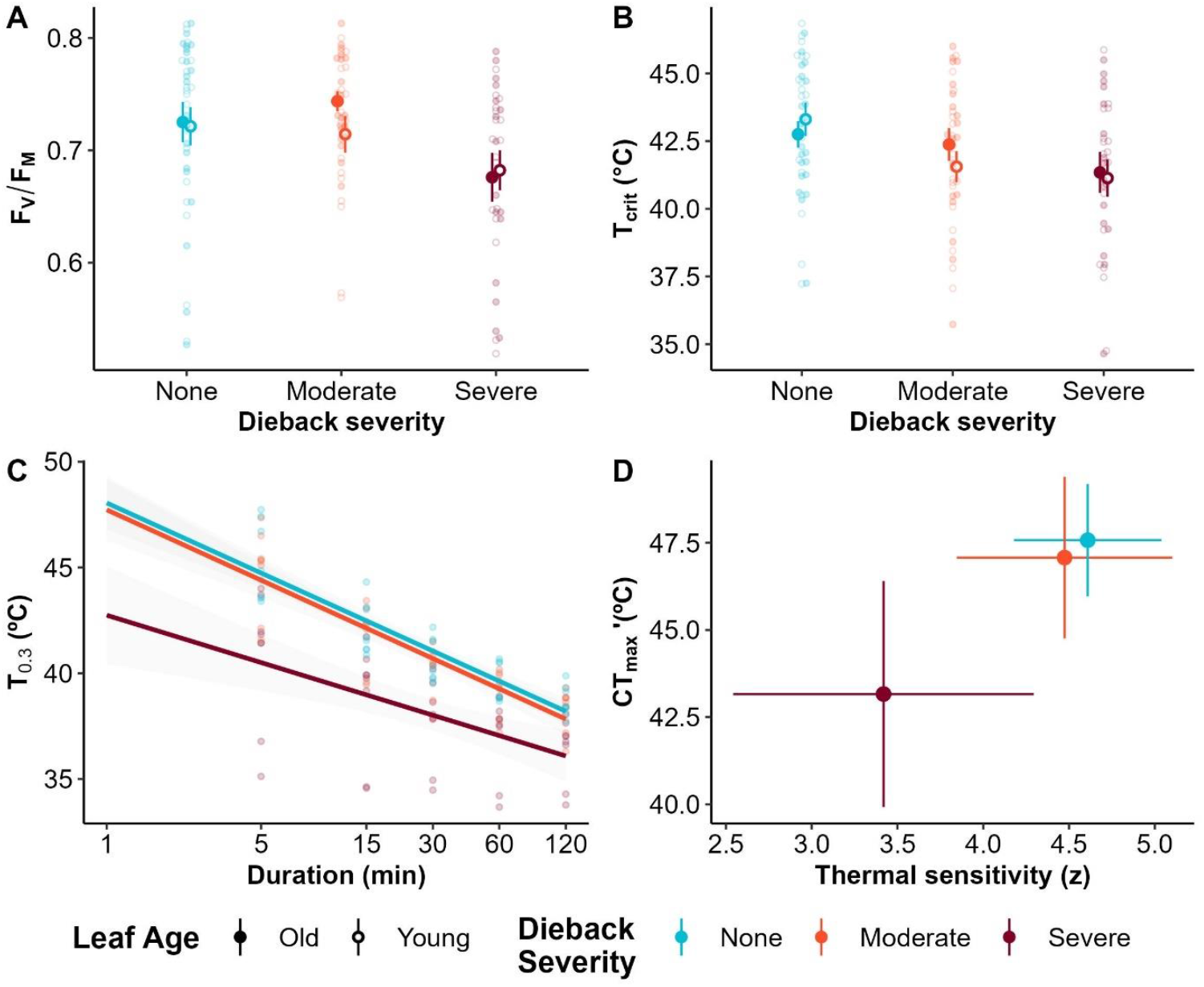
Changes in photosystem heat tolerance with increasing dieback severity in *Eucalyptus pauciflora* ssp. *niphophila*. **(A)** Pre-assay F_V_/F_M_ (maximum quantum yield of PSII); (**B)** T_crit_ (°C) (i.e. the upper temperature at which stress is incurred within PSII); **(C)** T_0.3_(°C) for any given duration (i.e. the temperature at which F_V_/F_M_ = 0.3, indicative of irreversible damage to PSII, shaded error bars reflect 95% confidence interval; **(D)** Comparison of slope (thermal sensitivity, *z*) and intercept (CT_max_′; heat tolerance at 1 minute duration) from plot (C), error bars reflect standard deviation about the mean.

The thermal load sensitivity experiment demonstrated that increased duration of exposure significantly reduced the temperature at which plants reached an F_V_/F_M_ value less than 0.3 (T_0.3_) (Figure 3C, Table 2). Trees with severe dieback also had significantly lower values for critical temperature at 1 minute (CT_max_′), than unaffected individuals (Figure 3D, Table 2). The slope (thermal sensitivity parameter, *z*) was also significantly lower in severely dieback affected trees than unaffected trees (Figure 3D, Table 2). For comparisons of CT_max_′ to T_crit_ see S2d and Figure S5.

Modelled probability of leaf survival based on simulated leaf temperatures indicates that even at current temperatures and with the benefit of evaporative cooling through open stomata, leaves from severely dieback affected trees had dramatically reduced probability of survival. If these trees were drought stressed and stomata had closed, the probability of leaf survival was even lower (Figure 4A, B). When a future climate warming scenario with an increase of 4°C is added to this simulation, the model indicates that leaves of these severely affected trees would accumulate lethal heat loads about halfway through the growing season under severe water stress or after two months with the benefit of evaporative cooling (Figure 4C).

**Figure 4:**
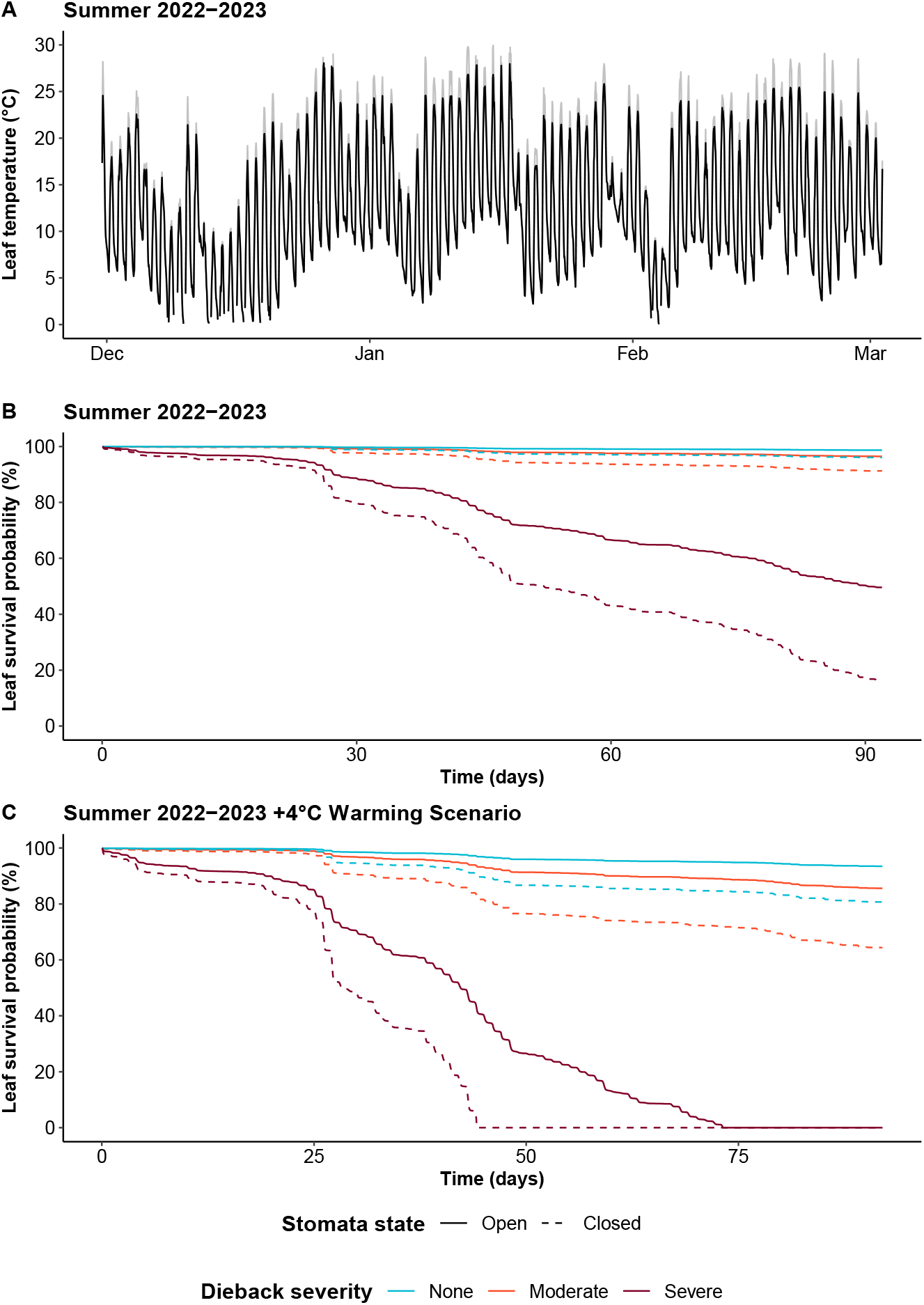
Modelled leaf temperatures and leaf survival probabilities across summer 2022-2023. **A)** Modelled leaf temperature for a standard *Eucalyptus pauciflora* ssp. *niphophila* leaf during the summer 2022-2023 growing season. Black lines indicate modelled leaf temperatures with open stomata, while light grey indicates leaf temperature with closed stomata. **B)** Declining leaf survival probability modelled based on cumulative heat load and TLS curve parameters for leaves of increasing dieback severity in the presence or absence of evaporative cooling (stomata open or closed). **C)** Leaf survival probability as per **B** but with a simulated +4°C increase in temperatures across the entire season.

## 4. Discussion

As global climate change places more tree species at risk of dieback, it is important to identify the drivers of dieback phenomena, and the physiology associated with decline or recovery of affected trees. This endeavour is complicated by challenges in distinguishing the causes from consequences of decline and often doesn’t incorporate feedback loops associated with trait and physiological tolerance changes in response to decline. In this study, we used the Australian snow gum, *E. pauciflora* spp. *niphophila*, as a case study to assess whether leaf traits relate to tree level vulnerability to the unfolding dieback event and explore how dieback-driven changes in functional traits may hasten tree decline. While we found significant changes in leaf traits in severely dieback affected trees (smaller leaves, higher stem to leaf area ratios, smaller stomata lengths and lower photosystem heat tolerance), the isolation of these changes to only the most severely affected individuals implies that these leaf traits likely do not contribute to initiation of woodborer infestation. Our findings also suggest that increased water stress, as evidenced by conservative leaf and branch traits in dieback affected snow gums, is accompanied by reduced vigour and physiological heat tolerance. Together with our models of accumulated damage from heat load our case study illustrates how these changes likely compound the direct hydraulic effects of woodborer damage through positive feedback cycles to limit carbon assimilation and growth, hastening insect-mediated decline.

### 4.1 Elevation effects on dieback driven by insect, not leaf traits

Our study was designed to enable separation of abiotic (elevation) effects and insect mediated effects. Accounting for spatial variation, we found no changes in leaf and branch morphology or stomatal anatomy along the abiotic elevation gradient, nor was there any interactive effect between elevation and dieback on leaf traits. Although *Eucalyptus pauciflora* ssp. *niphophila* trees growing at the lower edge of their elevational range were more frequently infested by *P. mastersi* than those at higher elevation sites, consistent with previous studies (Brookhouse et al., 2024; Bryant et al., 2024), the absence of an elevational effect on leaf traits suggests that the observed landscape patterning in snow gum dieback is more likely driven by biological and/or behavioural attributes of the insect, *P. mastersi*, rather than leaf traits of the affected trees. In the study of leaf, bark and hydraulic traits in *E. pauciflora* ssp. *niphophila* and closely related *E. pauciflora* ssp. *pauciflora* which occurs at lower elevations, Bryant et al. (2024) found that LMA and Huber values both increase with elevation and bark thickness of branches declines, but these were considered over two subspecies (*niphophila* and *pauciflora*) and a wider elevation range which likely explains why similar changes in LMA and Huber values were not seen in the present study. Further examination of leaf and bark traits across the wider species distribution, and of the ecology of *P. mastersi*, are warranted to better understand patterns of its infestation and predict future impacts on the two subspecies distributions.

### 4.2 Increasing dieback severity and water limitation in snow gums

Significant changes in leaf morphology were evident in trees with the most severe dieback symptoms, but not those with moderate dieback, indicating that the trait changes are responses to rather than drivers of vulnerability. Severely affected trees had smaller individual leaf areas and higher Huber values. This altered morphology is consistent with changes due to water stress; a likely consequence of damage to the xylem caused by *P. mastersi* larvae. Limited water availability is often associated with a reduction in leaf area, and thus less surface area through which water loss can occur (Leigh et al., 2017; Shao et al., 2008), but smaller leaf area can also reflect the effects of low turgor on leaf expansion (Coussement et al., 2021). Similarly, the observed increases in Huber values are the result of either reduced leaf production (number and/or size) or increased senescence, both of which would improve water status as they contribute to maintaining favourable water potential gradients and limit the risk of xylem cavitation in severely dieback affected trees (Carter & White, 2009). These results support the conclusion that the observed morphological changes are part of the trees’ response to late-stage dieback and woodborer damage to xylem rather than an intrinsic leaf trait of vulnerable trees.

Stomatal traits also changed in response to increasing dieback severity, consistent with water limitation. Trees with moderate and severe dieback had significantly smaller stomata than unaffected trees. Trees in the moderate dieback category showed evidence of early canopy decline and visible *P. mastersi* feeding galleries or puckering bark, suggesting that some xylem disruption had already occurred and the tree was likely operating with reduced canopy water supply. Smaller stomata in moderately and severely affected trees may therefore have arisen as a consequence of this water limitation as greater water limitation is generally associated with reduced stomatal size and density, reducing overall pore area and water loss (Bertolino et al., 2019; Driesen et al., 2023; Henry et al., 2019; Hetherington & Woodward, 2003; Zhao et al., 2015). Smaller stomata can also open and close more rapidly than larger stomata offering greater physiological stomatal control that may be advantageous in unfavourable conditions (Drake et al., 2013; Hetherington & Woodward, 2003; Raven, 2014). Despite having smaller stomata, dieback-affected trees showed no change in stomatal pore index, a proxy for maximum potential conductance per unit area that integrates both stomata density and length (Sack et al., 2003). Thus the change in stomata length was offset by small increases in stomatal density (Haworth et al., 2023). As trees with severe dieback had smaller stomata, smaller leaves and higher Huber values, the overall effect would be lower water demand and greater stomatal control, but at the cost of reduced potential carbon gain.

Future consideration of the stomatal thermal dynamics and thermoregulation, in conjunction with assays of carbon isotopes and non-structural carbohydrates in underground storage organs and stems of snow gums and other dieback affected species would assist in identifying the extent to which carbon assimilation and growth are limited by stomatal closure or carbon starvation (Farquhar et al., 2003). The approaches used here, in conjunction with consideration of dendrochronological series may assist to identify wood and bark trait variation and reveal other changes in growth preceding dieback as well as the relationship between growth changes and climatic factors such as annual temperatures and precipitation (Brookhouse & Bi, 2009).

### 4.3 Dieback reduces physiological tolerance of heat

In addition to morphological changes associated with *P. mastersi* infestation, we found a significant reduction in the photosystem heat tolerance of leaves from severely dieback-affected trees. This lowered tolerance threshold was consistent across both point and heat load tolerance measurements. While some studies find that mild to moderate water limitation can trigger increased heat tolerance (Cook et al., 2021; Havaux, 1992; Valladares & Pearcy, 1997), our results suggest that once the combination of heat and dieback induced (i.e. water limitation) stresses is severe enough it triggers activation of multiple stress response pathways that limit the resources available for photosystem function (Zhu et al., 2021).

Interestingly, this finding suggests dieback may create stress that exceeds the acclimatory potential of thermal tolerance. Prior studies indicating an acclimatory effect of water limitation considered temperate and desert species and attribute increased tolerance to acclimation of leaves to higher leaf temperatures. For example, Cook et al. (2021) found that leaves of drought-affected individuals reach higher, more stressful temperatures potentially due to stomatal closure and limited capacity for transpirational leaf cooling, but these leaves also had higher heat tolerance. We found no evidence of increased heat tolerance in dieback affected trees. This could mean that these sub-alpine trees are not subject to conditions in which water limitation leads to substantially higher leaf temperatures, although our models suggest that stomatal closure can increase leaf temperatures by 2-3°C (Figure 4A). However, we suspect that the stress caused by woodborer damage may be significantly greater than stress faced by trees in the aforementioned drought studies; for example, the shrubs measured by Cook et al. (2021) were water limited for four weeks prior to heat tolerance measurement but were otherwise healthy. Trees with severe dieback showed evidence of significant woodborer damage (feeding galleries) that has likely been limiting canopy water and nutrient supply, leading to reductions carbon assimilation for an entire growing season (or longer), leaving these trees in a precarious position with few resources to cope with additional stresses. Further study to assess the generality of these results for other broadleaf, evergreen, temperate forests is needed as we know little about the physiological consequences of dieback in those systems.

This study is one of few to apply an emerging measure of thermal tolerance, thermal load sensitivity (TLS), to plants to assess the interactive effects of time and intensity of heat (Arnold et al., 2025). Results from our TLS assays generally aligned with T_crit_ results but provide more nuanced insight. We found that duration of heat exposure (heat load) affected the occurrence of heat stress, with the thermal sensitivity of all dieback categories increased by extended exposure durations. While photosystem heat tolerance of all dieback categories declined with increasing duration, severely affected trees had a significantly lower critical temperature (CT_max_′) than unaffected trees, indicating a reduced tolerance for high temperatures. In these severely affected trees, exposure to 40°C (leaf temperature) for as little as 15 minutes could lead to significant damage to PSII (Figure 3C). Even in the relatively mild conditions of the Kosciuszko region, leaf temperatures can regularly exceed air temperatures by 10°C or more (Salisbury & Spomer, 1964). Moreover, even at temperatures between 35-°40 C, the Calvin cycle can be impaired and chloroplastic heat shock proteins upregulated (Wahid et al., 2007), indicating that the potential for accumulation of heat-induced damage in our trees with severe dieback is not trivial.

The dynamic models of heat load further indicate that even under relatively mild conditions, such as those experienced during the 2022/2023 growing season, leaves of severely dieback affected snow gums were likely to have reduced survival probability, particularly if they do not have the benefit of transpirational cooling. It is worth noting that the 2022/2023 summer season was mild compared to average current summer temperatures. These models therefore also present a stark illustration of how a hotter summer would impact and exacerbate dieback effects, leading to greater leaf mortality and reduced potential for the trees to grow or recover from insect infestation. This result may be a harbinger of the feedback associated with insect mediated decline in forest stands elsewhere in the world.

### 4.4 Conclusions

Around the world forest ecosystems are exhibiting dieback events that are likely to herald widespread species turnover. These changes are particularly impactful in systems, like the Australian sub-alpine, where the forest canopy is largely monospecific. Broadleaf evergreen mixed forests dominate the southern hemisphere and tropical systems more broadly but little is known about the dynamics of forest decline in these systems. Decline or loss of dominant canopy species will alter the ecological, hydrological and cultural values of the system, and in some cases – including snow gums – stands to impact state and national water-supply and power-generation systems. Using snow gums as a case study this project has illustrated an approach to separate biotic and abitioc drivers of dieback and to detect the extent to which variation among traits along a dieback severity gradient are drivers versus consequences of decline. For snow gum, leaf trait variation is not associated with variation in vulnerability to dieback within the affected subspecies, though previous studies indicate that bark traits may explain variation among subspecies. The present study also demonstrates how changes in traits in response to water stress likely form part of a positive feedback loop that hastens tree decline by reducing both capacity for photosynthetic assimilation and physiological tolerance of heat. Our results indicate that the impact of heat load associated damage on long lived leaves of evergreen species is a critical knowledge gap in our understanding of global forest decline.

## References

Adams, H. D., Guardiola-Claramonte, M., Barron-Gafford, G. A., Villegas, J. C., Breshears, D. D., Zou, C. B., Troch, P. A., & Huxman, T. E. (2009). Temperature sensitivity of drought-induced tree mortality portends increased regional die-off under global-change-type drought. Proceedings of the National Academy of Sciences of the United States of America, 106(17), 7063–7066. 10.1073/PNAS.0901438106/SUPPL_FILE/0901438106SI.PDF

Allen, C. D., Macalady, A. K., Chenchouni, H., Bachelet, D., McDowell, N., Vennetier, M., Kitzberger, T., Rigling, A., Breshears, D. D., Hogg, E. H. (Ted), Gonzalez, P., Fensham, R., Zhang, Z., Castro, J., Demidova, N., Lim, J. H., Allard, G., Running, S. W., Semerci, A., & Cobb, N. (2010). A global overview of drought and heat-induced tree mortality reveals emerging climate change risks for forests. Forest Ecology and Management, 259(4), 660–684. 10.1016/J.FORECO.2009.09.001

Anderegg, W. R. L., Hicke, J. A., Fisher, R. A., Allen, C. D., Aukema, J., Bentz, B., Hood, S., Lichstein, J. W., Macalady, A. K., Mcdowell, N., Pan, Y., Raffa, K., Sala, A., Shaw, J. D., Stephenson, N. L., Tague, C., & Zeppel, M. (2015). Tree mortality from drought, insects, and their interactions in a changing climate. New Phytologist, 208(3), 674–683. 10.1111/NPH.13477

Anderegg, W. R. L., Kane, J. M., & Anderegg, L. D. L. (2012). Consequences of widespread tree mortality triggered by drought and temperature stress. Nature Climate Change 2012 3:1, 3(1), 30–36. 10.1038/nclimate1635

Andrus, R. A., Peach, L. R., Cinquini, A. R., Mills, B., Yusi, J. T., Buhl, C., Fischer, M., Goodrich, B. A., Hulbert, J. M., Holz, A., Meddens, A. J. H., Moffett, K. B., Ramirez, A., & Adams, H. D. (2024). Canary in the forest?—Tree mortality and canopy dieback of western redcedar linked to drier and warmer summers. Journal of Biogeography, 51(1), 103–119. 10.1111/jbi.14732

Arnold, P. A., Briceño, V. F., Gowland, K. M., Catling, A. A., Bravo, L. A., & Nicotra, A. B. (2021). A high-throughput method for measuring critical thermal limits of. Functional Plant Biology.

Arnold, P. A., Noble, D. W. A., Nicotra, A. B., Kearney, M. R., Rezende, E. L., Andrew, S. C., Briceño, V. F., Buckley, L. B., Christian, K. A., Clusella-Trullas, S., Geange, S. R., Guja, L. K., Jiménez Robles, O., Kefford, B. J., Kellermann, V., Leigh, A., Marchin, R. M., Mokany, K., & Bennett, J. M. (2025). A Framework for Modelling Thermal Load Sensitivity Across Life. Global Change Biology, 31(7), e70315. 10.1111/gcb.70315

Banks, J. (1982). The use of dendrochronology in the Interpretation of Dynamics of the Snow Gum Forest. Australian National Univeristy.

Banks, J. (1989). A history of forest fire in the Australian Alps. The Scientific Significance of the Australian Alps. irst Fenner Conference on the Environment, Canberra.

Ben Rejeb, I., Pastor, V., & Mauch-Mani, B. (2014). Plant Responses to Simultaneous Biotic and Abiotic Stress: Molecular Mechanisms. Plants 2014, Vol. 3, Pages 458-475, 3(4), 458–475. 10.3390/PLANTS3040458

Bentz, B. J., Rgnire, J., Fettig, C. J., Hansen, E. M., Hayes, J. L., Hicke, J. A., Kelsey, R. G., Negron, J. F., & Seybold, S. J. (2010). Climate Change and Bark Beetles of the Western United States and Canada: Direct and Indirect Effects. BioScience, 60(8), 602–613. 10.1525/BIO.2010.60.8.6

Bertolino, L. T., Caine, R. S., & Gray, J. E. (2019). Impact of stomatal density and morphology on water-use efficiency in a changing world. Frontiers in Plant Science, 10, 225. 10.3389/FPLS.2019.00225/BIBTEX

Boyd, I. L., Freer-Smith, P. H., Gilligan, C. A., & Godfray, H. C. J. (2013). The consequence of tree pests and diseases for ecosystem services. Science, 342(6160). 10.1126/SCIENCE.1235773/SUPPL_FILE/BOYD-SM.PDF

Brookhouse, M., & Bi, H. (2009). Elevation-dependent climate sensitivity in Eucalyptus pauciflora Sieb. Ex Spreng. Trees, 23, 1309–1320. 10.1007/s00468-009-0372-6

Brookhouse, M., Farrow, R., Meyer, J., McDougall, K., Ward-Jones, J., & Wright, G. (2024). Elevation and Land Management Explain the Incidence and Severity of Phoracantha-Induced Decline within High-Elevation Eucalypt Woodlands. 10.2139/SSRN.4693230

Bryant, C., Ball, M. C., Borevitz, J., Brookhouse, M. T., Carle, H., Cunningham, P., Davey, M., Davies, J., Eason, A., Erskine, J. D., Fuenzalida, T. I., Grishin, D., Harris, R., Kriticos, J., Midson, A., Nicotra, A. B., Nshuti, A., Ward-Jones, J., Yau, Y., … Bothwell, H. (2024). Elevation-dependent patterns of borer-mediated snow-gum dieback are associated with subspecies’ trait differences and environmental variation. Austral Ecology, 49(3), e13508. 10.1111/aec.13508

Camarero, J. J., Gazol, A., Sangüesa-Barreda, G., Oliva, J., & Vicente-Serrano, S. M. (2015). To die or not to die: Early warnings of tree dieback in response to a severe drought. Journal of Ecology, 103(1), 44–57. 10.1111/1365-2745.12295

Carter, J. L., & White, D. A. (2009). Plasticity in the Huber value contributes to homeostasis in leaf water relations of a mallee Eucalypt with variation to groundwater depth. Tree Physiology, 29(11), 1407–1418. 10.1093/TREEPHYS/TPP076

Choat, B., Brodribb, T. J., Brodersen, C. R., Duursma, R. A., López, R., & Medlyn, B. E. (2018). Triggers of tree mortality under drought. Nature, 558(7711), 531–539. 10.1038/s41586-018-0240-x

Cook, A., Berry, N., Milner, K., & Leigh, A. (2021). Water availability influences thermal safety margins for leaves. Functional Ecology, 35(10), 2179–2189. 10.1111/1365-2435.13868

Cook, A., Rezende, E., Petrou, K., & Leigh, A. (2024). Beyond a single temperature threshold: Applying a cumulative thermal stress framework to plant heat tolerance. Ecology Letters, 27(3), e14416. 10.1111/ele.14416

Coussement, J. R., Villers, S. L. Y., Nelissen, H., Inzé, D., & Steppe, K. (2021). Turgor-time controls grass leaf elongation rate and duration under drought stress. Plant, Cell & Environment, 44(5), 1361–1378. 10.1111/PCE.13989

Da Conceição Caldeira, M., Fernandéz, V., Tomé, J., & Pereira, J. S. (2001). Positive effect of drought on longicorn borer larval survival and growth on eucalyptus trunks. Departmento de Engenharia Florestal, Instituto Superior de Agronomia, Tapada Da Ajud. 10.1051/forest

Drake, P. L., Froend, R. H., & Franks, P. J. (2013). Smaller, faster stomata: Scaling of stomatal size, rate of response, and stomatal conductance. Journal of Experimental Botany, 64(2), 495–505. 10.1093/JXB/ERS347

Driesen, E., De Proft, M., & Saeys, W. (2023). Drought stress triggers alterations of adaxial and abaxial stomatal development in basil leaves increasing water-use efficiency. Horticulture Research, 10(6). 10.1093/HR/UHAD075

Faber, A. H., Ørsted, M., & Ehlers, B. K. (2024). Application of the thermal death time model in predicting thermal damage accumulation in plants. Journal of Experimental Botany, 75(11), 3467–3482. 10.1093/jxb/erae096

Farquhar, G. D., Hubick, K. T., & Ehleringer, J. R. (2003). Carbon Isotope Discrimination and Photosynthesis. 10.1146/Annurev.Pp.40.060189.002443, 40(1), 503–537. 10.1146/ANNUREV.PP.40.060189.002443

Gobiet, A., Kotlarski, S., Beniston, M., Heinrich, G., Rajczak, J., & Stoffel, M. (2014). 21st century climate change in the European Alps—A review. Science of The Total Environment, 493, 1138–1151. 10.1016/J.SCITOTENV.2013.07.050

Hammond, W. M., Williams, A. P., Abatzoglou, J. T., Adams, H. D., Klein, T., López, R., Sáenz-Romero, C., Hartmann, H., Breshears, D. D., & Allen, C. D. (2022). Global field observations of tree die-off reveal hotter-drought fingerprint for Earth’s forests. Nature Communications 2022 13:1, 13(1), 1–11. 10.1038/s41467-022-29289-2

Hanks, L. M., Paine, T. D., & Milla∼, J. G. (1991). Mechanisms of Resistance in Eucalyptus Against Larvae of the Eucalyptus Longhorned Borer (Coleoptera: Cerambycidae). Environ. Entomol, 20(6), 1583–1588.

Hanks, L. M., Paine, T. D., Millar, J. G., Campbell, C. D., & Schuch, U. K. (1999). Water relations of host trees and resistance to the phloem-boring beetle Phoracantha semipunctata F. (Coleoptera: Cerambycidae). Oecologia, 119(3), 400–407. 10.1007/S004420050801/METRICS

Hartmann, H., Bastos, A., Das, A. J., Esquivel-Muelbert, A., Hammond, W. M., Martínez-Vilalta, J., Mcdowell, N. G., Powers, J. S., Pugh, T. A. M., Ruthrof, K. X., & Allen, C. D. (2022). Climate Change Risks to Global Forest Health: Emergence of Unexpected Events of Elevated Tree Mortality Worldwide. Annual Review of Plant Biology, 73(Volume 73, 2022), 673–702. 10.1146/ANNUREV-ARPLANT-102820-012804/1

Havaux, M. (1992). Stress Tolerance of Photosystem II in Vivo: Antagonistic Effects of Water, Heat, and Photoinhibition Stresses. Plant Physiology, 100(1), 424. 10.1104/PP.100.1.424

Haworth, M., Marino, G., Materassi, A., Raschi, A., Scutt, C. P., & Centritto, M. (2023). The functional significance of the stomatal size to density relationship: Interaction with atmospheric [CO2] and role in plant physiological behaviour. The Science of the Total Environment, 863. 10.1016/J.SCITOTENV.2022.160908

Henry, C., John, G. P., Pan, R., Bartlett, M. K., Fletcher, L. R., Scoffoni, C., & Sack, L. (2019). A stomatal safety-efficiency trade-off constrains responses to leaf dehydration. Nature Communications, 10(1). 10.1038/S41467-019-11006-1

Hersbach, H., Bell, B., Berrisford, P., Hirahara, S., Horányi, A., Muñoz-Sabater, J., Nicolas, J., Peubey, C., Radu, R., Schepers, D., Simmons, A., Soci, C., Abdalla, S., Abellan, X., Balsamo, G., Bechtold, P., Biavati, G., Bidlot, J., Bonavita, M., … Thépaut, J.-N. (2020). The ERA5 global reanalysis. Quarterly Journal of the Royal Meteorological Society, 146(730), 1999–2049. 10.1002/qj.3803

Hetherington, A. M., & Woodward, F. I. & (2003). The role of stomata in sensing and driving environmental change. https://www.nature.com/nature

Hosseini, A., Hosseini, S. M., & Linares, J. C. (2019). Linking morphological and ecophysiological leaf traits to canopy dieback in Persian oak trees from central Zagros. Journal of Forestry Research, 30(5), 1755–1764. 10.1007/S11676-018-0805-4/FIGURES/3

IPCC. (2022). Climate Change 2022: Impacts, Adaptation, and Vulnerability. Contribution of Working Group II to the Sixth Assessment Report of the Intergovernmental Panel on Climate Change [H.-O. Pörtner, D.C. Roberts, M. Tignor, E.S. Poloczanska, K. Mintenbeck, A. Alegría, M. Craig, S. Langsdorf, S. Löschke, V. Möller, A. Okem, B. Rama (eds.)]. Cambridge University Press. 10.59327/IPCC/AR6-9789291691647

Jaime, L., Batllori, E., & Lloret, F. (2024). Bark beetle outbreaks in coniferous forests: A review of climate change effects. European Journal of Forest Research, 143(1), 1–17. 10.1007/s10342-023-01623-3

Jurskis, V. (2005). Eucalypt decline in Australia, and a general concept of tree decline and dieback. Forest Ecology and Management, 215(1–3), 1–20. 10.1016/j.foreco.2005.04.026

Kearney, M. R., Gillingham, P. K., Bramer, I., Duffy, J. P., & Maclean, I. M. D. (2020). A method for computing hourly, historical, terrain-corrected microclimate anywhere on earth. Methods in Ecology and Evolution, 11(1), 38–43. 10.1111/2041-210X.13330

Kearney, M. R., & Leigh, A. (2024). Fast, accurate and accessible calculations of leaf temperature and its physiological consequences with NicheMapR. Methods in Ecology and Evolution, 15(9), 1516–1531. 10.1111/2041-210X.14373

Kearney, M. R., & Porter, W. P. (2017). NicheMapR – an R package for biophysical modelling: The microclimate model. Ecography, 40(5), 664–674. 10.1111/ecog.02360

Kearney, M. R., & Porter, W. P. (2020). NicheMapR – an R package for biophysical modelling: The ectotherm and Dynamic Energy Budget models. Ecography, 43(1), 85–96. 10.1111/ecog.04680

Klinges, D. H., Duffy, J. P., Kearney, M. R., & Maclean, I. M. D. (2022). mcera5: Driving microclimate models with ERA5 global gridded climate data. Methods in Ecology and Evolution, 13(7), 1402–1411. 10.1111/2041-210X.13877

Körner, Ch., & Cochrane, P. M. (1985). Stomatal responses and water relations of Eucalyptus pauciflora in summer along an elevational gradient. Oecologia, 66(3), 443–455. 10.1007/BF00378313

Leigh, A., Sevanto, S., Close, J. D., & Nicotra, A. B. (2017). The influence of leaf size and shape on leaf thermal dynamics: Does theory hold up under natural conditions? Plant Cell and Environment, 40(2), 237–248. 10.1111/pce.12857

Lin, Y., Kuang, L., Tang, S., Mou, Z., Phillips, O. L., Lambers, H., Liu, Z., Sardans, J., Peñuelas, J., Lai, Y., Lin, M., Chen, D., & Kuang, Y. (2021). Leaf traits from stomata to morphology are associated with climatic and edaphic variables for dominant tropical forest evergreen oaks. Journal of Plant Ecology, 14(6), 1115–1127. 10.1093/JPE/RTAB060

Manion, P. (1991). Tree disease concepts. Prentice Hall inc.

Martin, P. A., Newton, A. C., Cantarello, E., & Evans, P. (2015). Stand dieback and collapse in a temperate forest and its impact on forest structure and biodiversity. Forest Ecology and Management, 358, 130–138. 10.1016/J.FORECO.2015.08.033

McDowell, N., Pockman, W. T., Allen, C. D., Breshears, D. D., Cobb, N., Kolb, T., Plaut, J., Sperry, J., West, A., Williams, D. G., & Yepez, E. A. (2008). Mechanisms of plant survival and mortality during drought: Why do some plants survive while others succumb to drought? New Phytologist, 178(4), 719–739. 10.1111/j.1469-8137.2008.02436.x

Meir, P., Mencuccini, M., & Dewar, R. C. (2015). Drought-related tree mortality: Addressing the gaps in understanding and prediction. New Phytologist, 207(1), 28–33. 10.1111/nph.13382

Mitchell, R. J., Beaton, J. K., Bellamy, P. E., Broome, A., Chetcuti, J., Eaton, S., Ellis, C. J., Gimona, A., Harmer, R., Hester, A. J., Hewison, R. L., Hodgetts, N. G., Iason, G. R., Kerr, G., Littlewood, N. A., Newey, S., Potts, J. M., Pozsgai, G., Ray, D., … Woodward, S. (2014). Ash dieback in the UK: A review of the ecological and conservation implications and potential management options. Biological Conservation, 175, 95–109. 10.1016/J.BIOCON.2014.04.019

Mitton, J. B., & Ferrenberg, S. M. (2012). Mountain Pine Beetle Develops an Unprecedented Summer Generation in Response to Climate Warming. The American Naturalist, 179(5), E163–E171. 10.1086/665007

Mueller-Dombois, D. (1988). Towards a Unifying Theory for Stand-Level Dieback. In Forests of the World (Vol. 17, Issue 2, pp. 249–251). https://about.jstor.org/terms

Neuner, G., & Buchner, O. (2023). The dose makes the poison: The longer the heat lasts, the lower the temperature for functional impairment and damage. Environmental and Experimental Botany, 212, 105395. 10.1016/J.ENVEXPBOT.2023.105395

Niinemets, U. (2001). Global-Scale Climatic Controls of Leaf Dry Mass per Area, Density, and Thickness in Trees and Shrubs. Ecology, 82(2), 453. 10.2307/2679872

Pinheiro, J., Bates, D., & R Core Team. (2023). nlme: Linear and Nonlinear Mixed Effects Models [Computer software]. https://CRAN.R-project.org/package=nlme

Poorter, H., Niinemets, Ü., Poorter, L., Wright, I. J., & Villar, R. (2009). Causes and consequences of variation in leaf mass per area (LMA): A meta-analysis. New Phytologist, 182(3), 565–588. 10.1111/J.1469-8137.2009.02830.X

R Core Team. (2021). R: A language and environment for statistical computing [Computer software]. R Foundation for Statistical Computing.

Raffa, K. F., Aukema, B. H., Bentz, B. J., Carroll, A. L., Hicke, J. A., Turner, M. G., & Romme, W. H. (2008). Cross-scale Drivers of Natural Disturbances Prone to Anthropogenic Amplification: The Dynamics of Bark Beetle Eruptions. BioScience, 58(6), 501–517. 10.1641/B580607

Raven, J. A. (2014). Speedy small stomata? Journal of Experimental Botany, 65(6), 1415–1424. 10.1093/JXB/ERU032

Rezende, E. L., Bozinovic, F., Szilágyi, A., & Santos, M. (2020). Predicting temperature mortality and selection in natural Drosophila populations. Science, 369(6508), 1242–1245. 10.1126/science.aba9287

Rezende, E. L., Castañeda, L. E., & Santos, M. (2014). Tolerance landscapes in thermal ecology. Functional Ecology, 28(4), 799–809. 10.1111/1365-2435.12268

Ross, C., & Brack, C. (2015). Eucalyptus viminalis dieback in the Monaro region, NSW. Australian Forestry, 78(4), 243–253. 10.1080/00049158.2015.1076754

Sack, L., Cowan, P. D., Jaikumar, N., & Holbrook, N. M. (2003). The ‘hydrology’ of leaves: Coordination of structure and function in temperate woody species. Plant, Cell & Environment, 26(8), 1343–1356. 10.1046/J.0016-8025.2003.01058.X

Salisbury, F. B., & Spomer, G. G. (1964). LEAF TEMPERATURES OF ALPINE PLANTS IN THE FIELD. 497–505.

Schreiber, U., & Berry, J. A. (1977). Heat-induced changes of chlorophyll fluorescence in intact leaves correlated with damage of the photosynthetic apparatus. Planta, 136(3), 233–238. 10.1007/BF00385990/METRICS

Seaton, S., Matusick, G., Ruthrof, K. X., & Hardy, G. E. S. J. (2015). Outbreak of Phoracantha semipunctata in Response to Severe Drought in a Mediterranean Eucalyptus Forest. Forests 2015, Vol. 6, Pages 3868-3881, 6(11), 3868–3881. 10.3390/F6113868

Shao, H. B., Chu, L. Y., Jaleel, C. A., & Zhao, C. X. (2008). Water-deficit stress-induced anatomical changes in higher plants. Comptes Rendus Biologies, 331(3), 215–225. 10.1016/J.CRVI.2008.01.002

Shields, B. (1993). Patch dieback of the subalpine snow gum forest, E. pauciflora in Kosciusko National Park [BSc Honours]. Australian National Univeristy.

Valladares, F., & Pearcy, R. W. (1997). Interactions between water stress, sun-shade acclimation, heat tolerance and photoinhibition in the sclerophyll Heteromeles arbutifolia. Plant, Cell and Environment, 20(1), 25–36. 10.1046/J.1365-3040.1997.D01-8.X

Wahid, A., Gelani, S., Ashraf, M., & Foolad, M. R. (2007). Heat tolerance in plants: An overview. Environmental and Experimental Botany, 61(3), 199–223. 10.1016/j.envexpbot.2007.05.011

Wright, I. J., Dong, N., Maire, V., Prentice, I. C., Westoby, M., Díaz, S., Gallagher, R. V., Jacobs, B. F., Kooyman, R., Law, E. A., Leishman, M. R., Niinemets, Ü., Reich, P. B., Sack, L., Villar, R., Wang, H., & Wilf, P. (2017). Global climatic drivers of leaf size. Science, 357(6354), 917–921. 10.1126/SCIENCE.AAL4760/SUPPL_FILE/AAL4760-WRIGHT-SM_DATA_SET_S1.XLSX

Zhao, W., Sun, Y., Kjelgren, R., & Liu, X. (2015). Response of Stomatal Density and Bound Gas Exchange in Leaves of Maize to Soil Water Deficit. Acta Physiologiae Plantarum, 37(1). 10.1007/S11738-014-1704-8

Zhu, L., Wen, W., Thorpe, M. R., Hocart, C. H., & Song, X. (2021). Combining heat stress with preexisting drought exacerbated the effects on chlorophyll fluorescence rise kinetics in four contrasting plant species. International Journal of Molecular Sciences, 22(19), 10682. 10.3390/IJMS221910682/S1

