## Supplemental Information for "Morphological and heat-tolerance traits are associated with progression and impact of, but not vulnerability to, tree decline"

### Supporting Information: Morphological and heat-tolerance traits are associated with progression of, but not vulnerability to, tree decline in snow gum woodlands

#### S1. Species and site selection and field methods

This study was conducted in the Snowy River valley between 1680 and 1890 m asl in Kosciuszko National Park (KNP), New South Wales, Australia. The study area is characterised by a sub-alpine climate with cool summers and persistent snow-cover during winters (National Parks and Wildlife Service, 2003). Mean annual precipitation at the nearby Perisher Valley weather station is 1785 mm with most falls occurring during winter as snow. Mean monthly maximum temperatures peak in January at 19.6°C, with extreme maxima up to 30 °C, while mean monthly minima in winter reach -3.6°C during August (Australian Bureau of Meteorology, 2023). Consistent with forest composition throughout KNP above 1600 m asl, stands within the study area are comprised entirely of sparsely spaced *E. pauciflora* ssp. *niphophila* with a shrub or grassy understory. Overstorey trees have a multi-stemmed mallee growth form, with an average height of 10 m, in which many stems emerge from a subterranean lignotuber.

##### *S1a. Transect arrangement*

This study considered trees along 10 transects on the slopes of the mountain peak known as The Paralyser (1961 m) (Figure S1). These 10 transects represent a subset of 16 transects established for ongoing monitoring in collaboration with the NSW Department of Planning, Industry and Environment. Transects #1-8 extended from 1680 m asl to 1860 m asl and each had five study plots at elevations of 1680, 1710, 1760, 1810 and 1860 m asl. The lowest forested elevation at transects #11, 14 and 16 was 1760 m asl so only three or four plots were established on each of these transects at 1760, 1810, 1860 and 1890 m asl (Figure 1).

This transect arrangement was designed to cover a gradient of both elevation and dieback severity. Dieback in this area is most severe close to Guthega (transect 1) while sites closer to Kosciuszko Road (transect 16) remain largely unaffected (Figure 1). This design therefore maximized the ability to separate elevation-dependent variation in leaf traits from variations arising from dieback itself.

In total 46 study plots were considered. Variable radius plots (VRP) were established at each of these locations to document and compare the level of dieback severity. The VRP method was chosen as it allowed for a plot of 10-20 trees to be defined at all sites despite differing stand densities. A basal area wedge was used to assess which trees would be included in the plot. If a tree was borderline in the wedge it was included. If a tree had multiple stems each stem was individually measured and assessed. Diameter at breast height (DBH), height, canopy health and the extent of woodborer damage (frass-clearing holes, bark puckering and/or galleries) were recorded for all stems in the plot.

##### *S1b. Dieback severity and types of borer damage*

Dieback severity was scored on a scale from 0-4 (None-Severe) based on both the health of the canopy and the types of wood-borer damage present. The types of wood borer damage noted included frass-clearing holes, puckering bark and feeding galleries. Frass-clearing holes are small holes in bark through which the larvae living below pushes out frass (chewed up wood). These are the first visible signs of wood-borer infestation (Figure S1) and are usually found in horizontal or vertical lines and are accompanied by leaking red kino (sap) that stains surrounding bark. As larvae continue to feed below the bark, it begins to "pucker" or cave inwards into void below. This level of damage is usually associated with the beginning of canopy thinning. As the tree condition deteriorates further, the bark generally cracks and falls away from the tree revealing characteristic horizontal feeding galleries below. This is usually accompanied by significant loss of canopy (Figure S1).

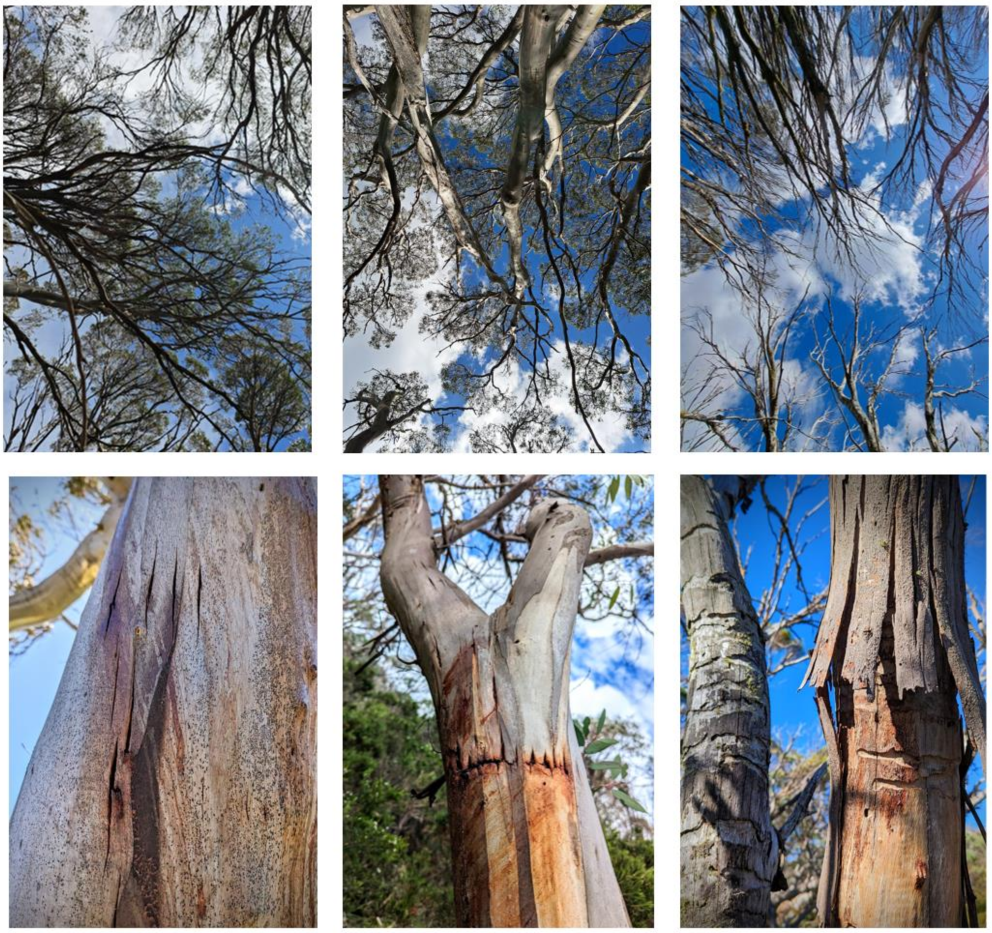

**Figure S1.** Increasing dieback severity in snow gum trees from sampled transects. Upper row is the canopy and lower is the stem of the same condition. Lefthand images show early signs of borer damage (frass-clearing holes) and fairly healthy canopy. Middle images show progressing borer damage with puckering bark, canopy thinning and some dead branches. Righthand images show late stages of borer damage with bark falling away from tree and little to no canopy remaining.

Canopy health was rated from 0-4 based on the following criterion: 4=Excellent, 75-100% of canopy healthy and intact, 3=Good, 50-75% canopy healthy and intact, 2=Fair, 25-50%, 1=Poor, <25% and 0=No canopy present. Woodborer damage was noted as the presence or absence of different types of damage including frass-clearing holes, bark puckering and characteristic horizontal feeding galleries. These features were chosen as they signify differing levels of dieback, i.e. frass-clearing holes indicate recent infestation whereas galleries indicate longer term infestation and more damage to the tree vascular system (Brookhouse et al., 2024). For the present study, these observations of canopy health and borer damage were used to assign a dieback severity rating on a four point scale where: "None" indicates no signs of woodborer damage, "Low" applies to trees with frass-clearing holes or minor bark puckering, and canopy health score of 3 or 4, "Moderate" means there is bark puckering and/or visible feeding galleries, canopy health score is 3 and "Severe" for trees with bark puckering and/or visible feeding galleries, canopy health score of 1 or 2.

##### *S1c. Distribution of dieback across the study site*

This study sampled leaves from 138 trees across 10 transects and at elevations from 1680 m to 1890 m asl. The prevalence of dieback differed with both elevation and transect number. As expected, dieback was more severe at transects #1-7 (those transects closer to Guthega), compared to transects #11, 14, 16 (those transects closer to Kosciuszko Road) (Figure S2A). Dieback was also more severe at the lowest elevations (1680 m and 1710 m). About 50% of the trees sampled at these elevations had severe levels of dieback and there was only one sampled tree that had no evidence of dieback (Figure S2B). At 1760 m most trees had low or moderate levels of dieback and above this elevation most trees were unaffected (Figure S2B).

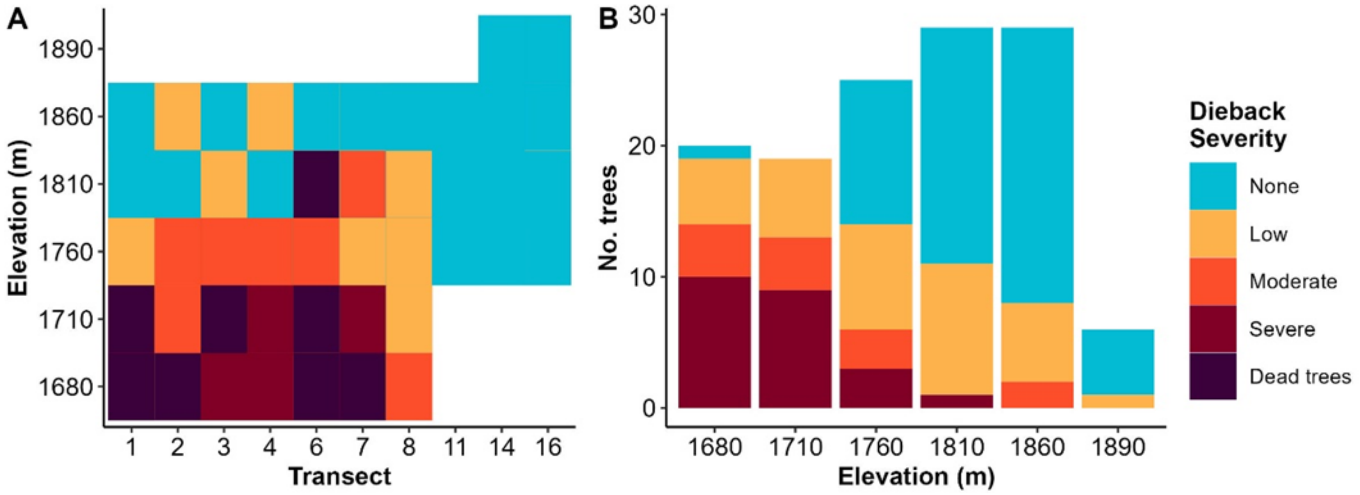

**Figure S2**. The distribution of dieback affected trees across elevation and transects. **(A)** The modal dieback severity score among all trees (n=7 to n=20) in each variable radius plot; **(B)** The number of sampled trees at each elevation within each dieback severity class.

##### *S1d. Classification of old and young leaves*

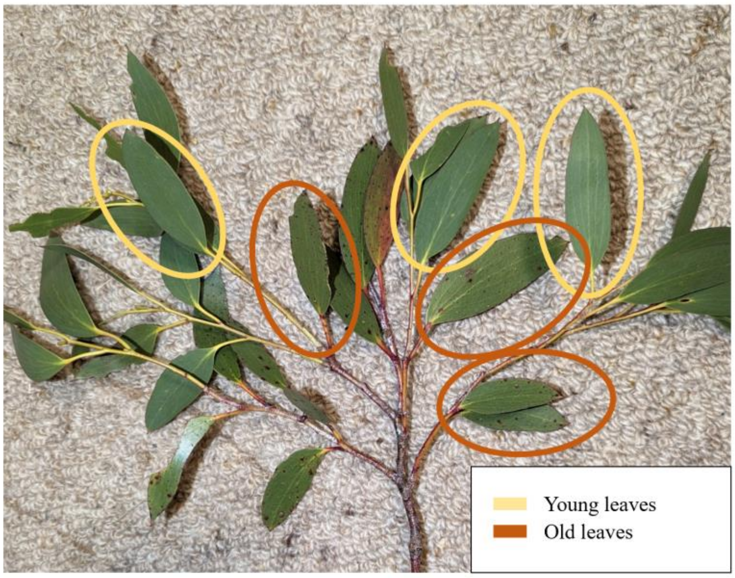

**Figure S3.** Example of branch with both young (expanded in 2022/23 growing season) and old (expanded in 2021/22 growing season) leaves. Young leaves were always located furthest from the main branch and were lighter in colour with pale green petioles. Old leaves were usually darker green and had red petioles.

##### *S1e. Leaf and stem trait measurement*

The area of each sampled leaf was measured using the Leaf Scan app (Anderson & Rosas-Anderson, 2017). Leaves were then oven-dried for 48 hours at 60 °C and weighed to obtain a dry mass and calculate leaf mass per unit area (LMA, g $cm^{-2}$). Huber values were calculated for one small branch (diameter approx. 5 mm) per tree. Digital callipers were used to measure stem diameter at the base of the branch, then the number of leaves of each leaf age were counted for later calculation of total leaf area using the average area of leaves from corresponding trees.

Stomatal impressions were collected from one leaf of each leaf age from each tree by applying clear nail polish to a 20 × 10 mm area. Nail polish was applied midway between mid-vein and the leaf margin. After allowing the polish to dry, it was peeled away from the leaf and the impression it held was photographed under an optical microscope at ×40 magnification with three different fields of view (FOV). Image analysis software 'ImageJ' (Schindelin et al., 2012) was used to measure guard-cell length (SL) and stomatal density per unit area (SD). The length of six stomata were measured for each leaf, two from each of three FOVs and the mean of these six calculated. Density was calculated by counting all stomata (where at least 50% of the stomatal cell was in the FOV) in each of the three FOVs and these values were averaged for each leaf. Stomatal pore index (SPI) was calculated as a description of total conductance potential, considering both stomatal size and density (Lin et al., 2021; Sack et al., 2003), expressed as:

Eq. 1 $SPI = SD \times SL^2 \times 10^{-4}$

#### *S1f. Notes on sample size*

Leaves were sampled from a total of 128 trees giving a maximum leaf sample size of 128 trees x 2 leaf ages x 3 replicates = 768 leaves. Not all trees had sufficient leaves or new leaves so total sample was slightly smaller. See Table S1 and S2 for further breakdown of sample sizes.

**Table S1**. Distribution of sampled leaves across dieback classes and notes where samples are missing. See Table S2. for outline of reasons for missing samples.

| | **Leaf area, LMA** | | **Huber value** | | **Stomata length, density** | |
|---|---|---|---|---|---|---|
| | **Max** | **Actual** | **Max** | **Actual** | **Max** | **Actual** |
| None - Old | 171 | 164 | 57 | 48 | 57 | 51 |
| None - Young | 171 | 171 | | | 57 | 50 |
| Low - Old | 114 | 111 | 38 | 35 | 38 | 35 |
| Low - Young | 114 | 114 | | | 38 | 37 |
| Moderate - Old | 36 | 36 | 12 | 12 | 12 | 11 |
| Moderate - Young | 36 | 36 | | | 12 | 12 |
| Severe - Old | 63 | 59 | 21 | 19 | 21 | 18 |
| Severe - Young | 63 | 57 | | | 21 | 19 |
| **Total** | 768 | 748 | 128 | 114 | 256 | 233 |

**Table S2.** The maximum sample size for each group of leaf/branch traits based on sampling 128 trees and notes why actual analysed sample sizes diverged from these maximum values.

| | **Leaf area, LMA** | **Huber value** | **Stomata length, density** |
|---|---|---|---|
| **Maximum sample size** | 768 | 128 | 256 |
| **Reason for missing sample:** | | | |
| Branches sampled, but contained no old leaves | -14 | | -4 |
| Branches sampled, but contained no young leaves | -6 | | -2 |
| Sample branch was not large enough to properly measure stem diameter for HV (most of these were from our first day sampling after which larger branches were collected) | | -14 | |
| Failed stomata peels, no photo available for ImageJ analysis | | | -3 |
| Blurry stomata pic, could not properly discern edge of stomata cells so excluded from ImageJ analysis | | | -10 |
| Excluded from stomata analysis due to being overly influential (as the only tree for certain dieback class at certain elevation) | | | -4 |
| **Actual sample size** | 748 | 114 | 233 |

#### S2. Photosystem Thermal Tolerance Methods

For both heat tolerance assays, leaves were randomly selected from sampled branches and cut into ~20 × 20 mm segments using a razor blade. Leaves with significant discolouration or herbivory were avoided. Young (2022/23) and old (2021/22) leaves were separated to test for an effect of leaf age on heat tolerance. Once cut, leaf segments were stored in labelled zip-lock bags on moist paper towel to prevent desiccation until test arrays were constructed (usually within 2 hours). To assemble test arrays, leaf segments were randomly assigned a grid location on a 72-square paper array and attached using double-sided tape.

##### *S2a. $T_{crit}$ assay*

The maximum quantum yield of photosystem II (PSII) ($F_V/F_M$) was measured for each leaf on the 72-square array prior to any heat treatment using a Maxi-Imaging PAM (Pulse-Amplitude Modulated chlorophyll-fluorescence imaging system; Heinz Walz GmbH, Effeltrich, Germany), to provide a rapid measurement of initial leaf photosystem health. As per Arnold et al. (2021), arrays of leaves were dark-adapted for 30 minutes prior to measurement of $F_0$; the fluorescence emitted by a dark-adapted leaf when exposed to a low intensity measuring light at 1 Hz. The arrays were then exposed to a saturating pulse to determine the maximal fluorescence ($F_M$) when the photosystem reaction centres are closed. Variable fluorescence ($F_V$) was calculated as $F_M – F_0$ and the relative maximum quantum yield of PSII photochemistry ($F_V/F_M$) was derived.

Following the measurement of $F_V/F_M$, we calculated $T_{crit}$ for each leaf segment on the 72-square array by assessing the temperature-dependent change in basal chlorophyll fluorescence ($F_0$) with increasing temperature (T). Basal chlorophyll fluorescence was continually measured using the Maxi-Imaging PAM as leaf temperature was ramped from 20°C to 70°C at a rate of 30°C $h^{-1}$ using a thermoelectric Peltier plate (Arnold et al., 2021). Throughout the experiment, leaf temperature was recorded using fine-wire type-T thermocouples (36-gauge, Omega Engineering, Norwalk, CT, USA) and a datalogger (DataTaker DT85; Lontek, Australia). Using the change in $F_0$ as leaf temperature increased, a T-$F_0$ curve was plotted for each leaf segment. We used breakpoint regression analysis with the *segmented* package in R (Muggeo, 2017) to define $T_{crit}$.

##### *S2b. Thermal load sensitivity of photosystem II*

For the TLS protocol, we used the same 72-square array as above and subjected arrays to one of seven temperatures (25, 34, 37, 40, 43, 46, and 49°C) and five durations (5, 15, 30, 60 and 120 minutes). For heat treatment each array was sealed in a transparent plastic bag and fully submerged in a water bath. Water baths were heated using SousVide Precision Cookers (Kogan, VIC, Australia). Fine-wire thermocouples and a datalogger were used to monitor water bath temperature, which remained within ± 0.5 °C of the set temperature for the duration of the treatment. A thin wire rack was used to keep each array flat and submerged.

To commence the treatment sequence, arrays were dark-adapted for 30 minutes before an initial $F_V/F_M$ measurement was taken using the Maxi-Imaging PAM. Each array was then light-adapted for 15 minutes under a 15 W LED grow light (~300 µmol $m^{-2}$ $s^{-1}$ PPFD). Arrays were consistently exposed to this sub-saturating light level for the duration of the heat treatment as previous studies identified light as an essential modulator of damage during heat stress (Curtis et al., 2014). Following the period of light-adapting, arrays were subjected to a randomly assigned temperature × duration treatment. Following the treatment all arrays were returned to the dark and left overnight before a final $F_V/F_M$ measurement was taken the next day, approximately 16 h post-treatment, which has been shown to allow for potential recovery or expression of damage (Curtis et al., 2014). These $F_V/F_M$ values were then used to plot a TLS curve that identifies the temperature at which loss of function of PSII occurs for any given duration (Cook et al., 2024; Rezende et al., 2014). Following Cook et al., (2024), boot-strapped regression over 1000 iterations was used to fit logistic curves to measured $F_V/F_M$ values to model decay of $F_V/F_M$ with increasing temperature and duration (see Figure S4). The temperature at which $F_V/F_M = 0.3$ (hereafter $T_{0.3}$) was extracted from each duration curve. We selected this as a threshold $F_V/F_M$ value at which significant damage has occurred to PSII which is irreversible over the 16-hours period (Curtis et al., 2014). While it is more common to use $T_{50}$, the point at which $F_V/F_M$ has declined by 50% from an initial point (healthy leaves having initial $F_V/F_M$ of 0.7-0.85), we considered the absolute measure $F_V/F_M$ of 0.3 to be a more standardised indication of irreversible damage as it was not influenced by the $F_V/F_M$ of the leaves prior to heat treatment.

To compare the heat tolerance of trees with differing dieback severity, $T_{0.3}$ was plotted as a response to heat exposure duration for each sampled tree and linear models were fitted separately for each level of dieback severity. $T_{0.3}$ was $\log_{10}$ transformed to allow for the fitting of this linear model which was then extrapolated to a duration of 1 minute to estimate the maximum heat tolerance (heat tolerance at 1 minute; $CT_{max}'$) (Cook et al., 2024; Rezende et al., 2014). The slope of this linear model defined the thermal sensitivity parameter ($z$), and this was also compared across dieback severities. Because preliminary analysis indicated that there was no influence of leaf age on heat tolerance, leaf age was not included in the modelling.

##### *S2c. Thermal load sensitivity model curve fitting*

In analysis of heat load data, logistic regression models were fitted to measured $F_V/F_M$ data to examine the decline of $F_V/F_M$ with temperature and duration combinations (Figure S4). There was a steeper decline in $F_V/F_M$ with temperature with increasing duration for all three dieback severities (Figure S4). $F_V/F_M$ also decayed in a logarithmic manner with increasing stress duration for the four highest temperature treatments (40, 43, 46, 49°C) (Figure S4). There was little to no change in $F_V/F_M$ with duration for temperatures below 40°C (25, 34, 37°C) (Figure S4B,D,F). These curves were plotted using relative $F_V/F_M$ data (scaled 0-1) to illustrate changes in the shape of the decay curve with duration and temperature. Final analysis and calculated tolerance thresholds used unscaled $F_V/F_M$ data.

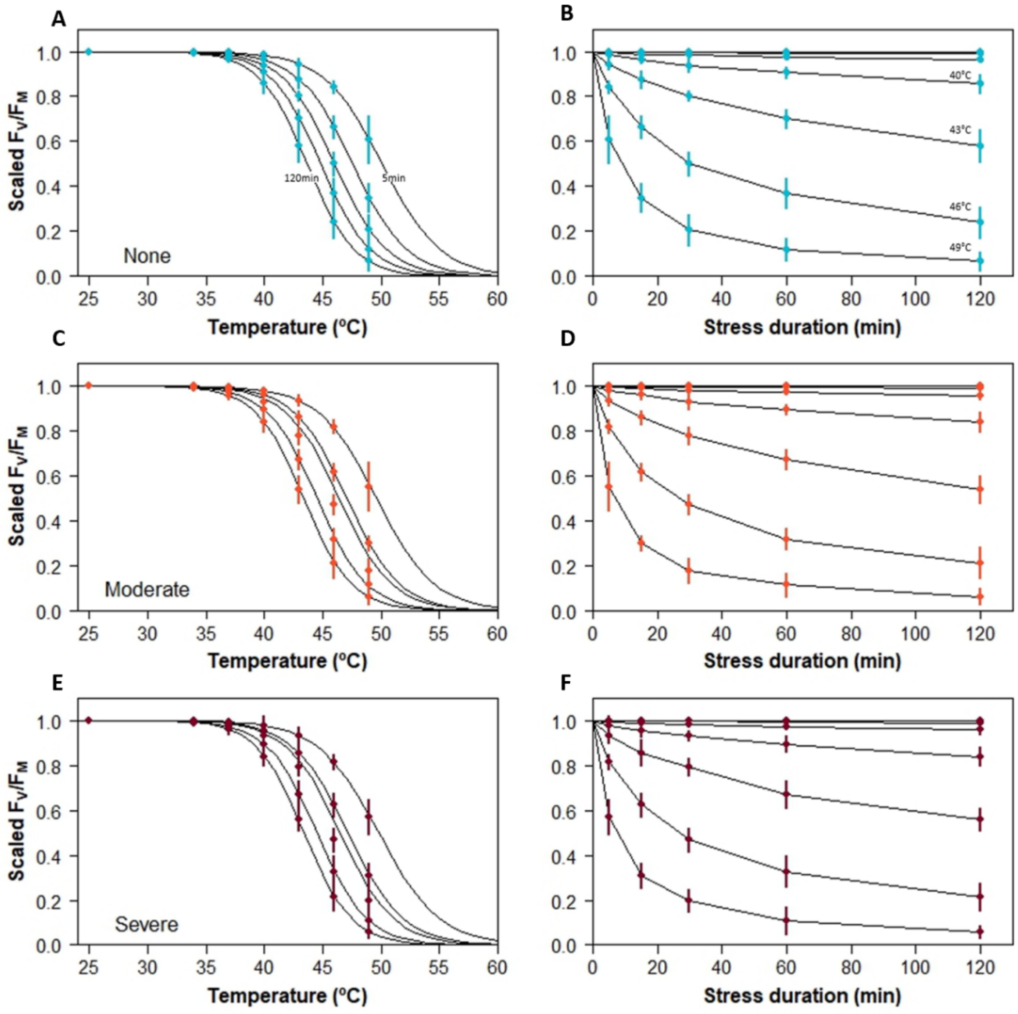

**Figure S4.** Decline in scaled $F_V/F_M$ with treatment temperature and stress duration in *Eucalyptus pauciflora* ssp. *niphophila*. Left hand panels show fitted logistic models with one line per duration (5, 15, 30, 60, 120 minutes). Right hand panels plot model predicted scaled $F_V/F_M$ with increasing duration, each line represents a different temperature treatment (25, 34, 37, 40, 43, 46, 49°C). Error bars reflect standard deviation in model estimate.

##### *S2d. Comparisons of photosystem heat tolerance metrics*

Trees with higher initial maximum quantum yield of PSII ($F_V/F_M$) generally had a higher $T_{crit}$ (Figure S5A). The calculation of heat tolerance parameters under heat load (exposure duration) is only recently being measured, it was of interest to compare this outcome to $T_{crit}$, a more widely reported metric in plant ecological studies. The $T_{crit}$ threshold is reached at lower temperatures than the estimated $CT_{max}'$ (heat tolerance at 1 minute exposure) (Figure S5B). The relationship between these two is also not 1:1, rather the slope is shallower indicating that a rise of 1°C in $CT_{max}'$ would correspond to less than 1°C increase in $T_{crit}$ (Figure S5B).

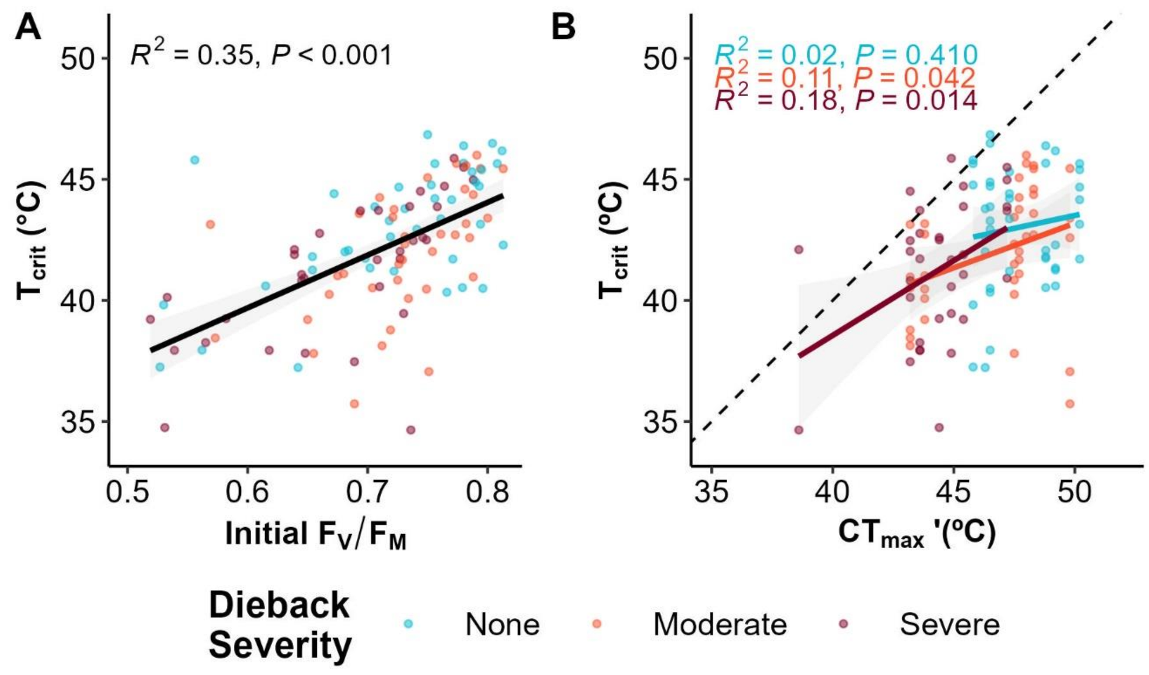

**Figure S5.** Comparisons of photosystem health and photosystem heat tolerance measures. **(A)** Linear correlation between increasing initial $F_V/F_M$ (i.e. maximum quantum yield of PSII) and $T_{crit}$ (i.e. the upper temperature at which stress is incurred in PSII) values; **(B)** Regression of $T_{crit}$ against $CT_{max}'$ (i.e. heat tolerance at 1 minute as derived from cumulative heat tolerance measurements).

#### S3. Data analysis supplement

Model comparison using the corrected Akaike Information Criterion (AICc) was undertaken to determine whether interaction terms improved model fit and therefore should be included in analysis. The R package 'AICcmodavg' (Mazerolle, 2023) was used to collate and compare AICc values for each model. For each leaf trait, models with no interaction, each possible 2-way interaction (leaf age x dieback, leaf age x elevation and elevation x dieback) and the 3-way interaction were compared. The "best" (i.e. most parsimonious) model based on this method was the model with the lowest AICc that differed from its counterparts by at least two points. For all traits, the most parsimonious model was that without any interaction terms, except for LMA where the selected model included the two-way interaction between leaf age and dieback (Table S3).

The R package ‘nlme’ offers five different forms of spatial correlation structures (linear, exponential, rational quadratic, spherical, and logistic). Models were created with each possible correlation structure and model AICc were again used to choose the most suitable structure, in this case the gaussian correlation structure (Table S4). This structure was included in the analyses of all transect-based variables.

**Table S3**. Model comparison using the corrected Akaike Information Criterion (AICc) to determine whether interaction terms improve the model fit. Models that were selected for interpretation were those with the lowest AICc and are indicated in bold.

\

| Leaf trait | No interactions | Leaf age x Dieback | Leaf age x Elevation | Elevation x Dieback | 3-way interaction |
|---|---|---|---|---|---|
| Leaf Area | **-33.61** | -15.42 | -17.01 | 7.89 | 83.98 |
| LMA | 7083.8 | **7059.7** | 7086.6 | 7094.8 | 7085.3 |
| Huber value | **570.1** | - | - | 590.8 | - |
| Stomata density | **-211.31** | -192.72 | -151.46 | -168.51 | -91.76 |
| Stomata length | **1230.9** | 1237.7 | 1296.8 | 1248.1 | 1276.7 |
| SPI | **-18.17** | -3.04 | 41.55 | 21.98 | 91.15 |
| $F_V/F_M$ | **-245.6** | -230.4 | - | - | - |
| $T_{crit}$ | **543.2** | 542.4 | - | - | - |

**Table S4** AICc values for linear mixed effects regression models with different spatial autocorrelation structures. The model AICc scores shown below were from the model of leaf area, but the gaussian correlation structure consistently performed best for all morphological leaf traits.

| Correlation structure | df | Model AICc |
|---|---|---|
| None | 7 | 23.87 |
| Exponential | 9 | -29.23 |
| **Gaussian** | **9** | **-33.91** |
| Linear | 9 | -33.57 |
| Rational quadratic | 9 | -32.68 |
| Spherical | 9 | -33.80 |

#### S4. Supplemental references

Anderson, C., & Rosas-Anderson, P. (2017). *Leafscan app* [Mobile application software]. https://www.leafscanapp.com

Arnold, P. A., Briceño, V. F., Gowland, K. M., Catling, A. A., Bravo, L. A., & Nicotra, A. B. (2021). A high-throughput method for measuring critical thermal limits of. *Functional Plant Biology*.

Brookhouse, M., Farrow, R., Meyer, J., McDougall, K., Ward-Jones, J., & Wright, G. (2024). *Elevation and Land Management Explain the Incidence and Severity of Phoracantha-Induced Decline within High-Elevation Eucalypt Woodlands*. https://doi.org/10.2139/SSRN.4693230

Cook, A., Rezende, E., Petrou, K., & Leigh, A. (2024). Beyond a single temperature threshold: Applying a cumulative thermal stress framework to plant heat tolerance. *Ecology Letters*, *27*(3), e14416. https://doi.org/10.1111/ele.14416

Curtis, E. M., Knight, C. A., Petrou, K., & Leigh, A. (2014). A comparative analysis of photosynthetic recovery from thermal stress: A desert plant case study. *Oecologia*, *175*(4), 1051–1061. https://doi.org/10.1007/s00442-014-2988-5

Lin, Y., Kuang, L., Tang, S., Mou, Z., Phillips, O. L., Lambers, H., Liu, Z., Sardans, J., Peñuelas, J., Lai, Y., Lin, M., Chen, D., & Kuang, Y. (2021). Leaf traits from stomata to morphology are associated with climatic and edaphic variables for dominant tropical forest evergreen oaks. *Journal of Plant Ecology*, *14*(6), 1115–1127. https://doi.org/10.1093/JPE/RTAB060

Mazerolle, M. (2023). *AICcmodavg: Model selection and multimodel inference based on (Q)AIC(c)* [Computer software]. https://cran.r-project.org/package=AICcmodavg

Muggeo, V. (2017). Interval estimation for the breakpoint in segmented regression: A smoothed score-based approach. *Australian & New Zealand Journal of Statistics*, *59*(3), 311–322.

National Parks and Wildlife Service. (2003). *Bioregions of NSW - The Australian Alps Bioregion*.

Rezende, E. L., Castañeda, L. E., & Santos, M. (2014). Tolerance landscapes in thermal ecology. *Functional Ecology*, *28*(4), 799–809. https://doi.org/10.1111/1365-2435.12268

Sack, L., Cowan, P. D., Jaikumar, N., & Holbrook, N. M. (2003). The 'hydrology' of leaves: Co-ordination of structure and function in temperate woody species. *Plant, Cell & Environment*, *26*(8), 1343–1356. https://doi.org/10.1046/J.0016-8025.2003.01058.X

Schindelin, J., Arganda-Carreras, I., Frise, E., Kaynig, V., Longair, M., Pietzsch, T., Preibisch, S., Rueden, C., Saalfeld, S., Schmid, B., Tinevez, J.-Y., White, D. J., Hartenstein, V., Eliceiri, K., Tomancak, P., & Cardona, A. (2012). Fiji: An open-source platform for biological-image analysis. *Nature Methods*, *9*(7), 676–682. https://doi.org/10.1038/nmeth.2019
